# Immune-mediated Engagement of T Regulatory Cells with Tumor Cells Results in Trogocytosis and Tumor Cell Killing

**DOI:** 10.64898/2025.12.11.693807

**Authors:** Amy K Erbe, Arika S. Feils, Anqi Gao, Sabrina N. VandenHeuvel, Simon Boyenga, Alina Hampton, Mackenzie Heck, Jen Zaborek, Daniel Spiegelman, Noah Tsarovsky, Anna Hoefges, Alexander A. Pieper, Peter M. Carlson, Mildred Felder, Ravi B Patel, Stephen D. Gillies, Alexa Heaton, Zachary Morris, Huy Q. Dinh, Alexander L. Rakhmilevich, Paul M. Sondel

**Author notes:** **<u>Corresponding Author:</u>** Amy Erbe PhD Research Asst. Professor University of Wisconsin, UW Carbone Cancer Center.

## Abstract

Despite advances in treatment, >50% of patients with advanced melanoma are unresponsive to current therapies. Using the B78-D14 melanoma model (GD2^+^/MHC-I⁻/MHC-II^+^), we can cure mice with a regimen that includes radiation therapy (RT) in combination with immunocytokine (IC; anti-GD2 monoclonal antibody linked to IL-2) while establishing immunological memory. We interrogated the role of T cells in the antitumor and memory responses following RT+IC. We show a requirement for CD4, but not CD8 T cells, to achieve both the initial and memory responses. Upon IC-induced cell-cell contact, subsets of CD4 T cells, including Foxp3⁺ T regulatory cells, trogocytose GD2 from tumor cells, acquire cytotoxic granules, and kill tumor cells. These results were confirmed using human tumor cell lines. These findings reveal that CD4⁺ T regulatory cells, upon immunologically-induced binding to tumor cells, can trogocytose tumor antigens and directly kill tumor cells, redefining their potential role in antitumor immunity.

## Introduction

The incidence of melanoma in the U.S. continues to rise. Among patients with advanced disease, the current standard of care—immune checkpoint inhibition (ICI)—achieves a 5-year progression-free survival (PFS) rate of less than 30%.^1,2^ Tumors that fail to respond to ICI often harbor features that promote growth while suppressing or evading immune recognition and elimination.^3–5^ Although ICI has markedly improved outcomes across several cancers, including melanoma, its benefit is not universal.^4,6^ Melanomas with a high tumor mutation burden (TMB) are considered “hot” and have actionable mutation-driven neoantigens, prominent Major Histocompatibility Complex class I (MHC-I) expression, and substantial T cell infiltration prior to therapy. These characteristics indicate an active endogenous immune response that is being dampened by checkpoint signaling. Patients with “hot” melanomas are more likely to benefit from ICI;^7–9^ however >70% of advanced melanoma patients do not show prolonged PFS in response to ICI. Most of these non-responsive tumors are considered “cold”, characterized by a low actionable TMB, poor immune infiltration, and/or loss of MHC-I.^10–15^

T cells can be primed to specifically recognize, remember, and eliminate melanoma by engaging melanoma-derived peptides presented in the cleft of MHC-I and/or MHC-II molecules on tumor cells. The resulting immune activation and memory enable ongoing surveillance, through which T cells identify and eradicate refractory, recurrent, or disseminated melanoma cells unless their activity is suppressed by immune checkpoints—circumstances under which ICI therapy can restore function. However, downregulation or loss of MHC-I is common in melanoma, preventing recognition by tumor-specific CD8 T cells (CD8s). Thus, MHC-I loss is a common mechanism of immune evasion and resistance to therapy.^5,16,17^ Separately, MHC-II molecules can present antigen to CD4 T cells (CD4s), initiating a cascade of immune responses. While not commonly expressed on most solid tumors and usually restricted to expression on immune cells, MHC-II is expressed on 50-60% of melanomas, but the implications of MHC-II expression on melanoma are not well understood.^16,18–21^ Whether MHC-II^+^ tumors directly, or indirectly, engage CD4s likely depends on the tumor microenvironment (TME) and treatment.^22^ CD4 effector, helper, and T regulatory (Treg) cells can be activated through MHC-II. Within the TME, Tregs may expand preferentially in MHC-II⁺ tumors, and stimulation of Tregs through MHC-II interactions can enhance their immunosuppressive activity.^23^ Yet paradoxically, patients with MHC-II⁺ melanomas tend to respond more favorably to ICI therapy.^21^ Thus, it is possible the antitumor response to ICI for patients with MHC-II^+^ tumors may reflect a balance of Treg versus T effector (T_eff_) activation prior to ICI treatment, though additional data are needed to clarify these mechanisms.

To investigate how MHC-I/II expression influences T cell responses induced by immunotherapy, we used the “immunologically cold” B78-D14 murine melanoma tumor model, which is poorly infiltrated by immune cells and resistant to ICI alone.^24–27^ B78-D14 cells (B78-D14s) express GD2, a tumor-antigen expressed on many melanomas, but expressed minimally on most normal tissues. Using this translationally relevant model, combining radiation therapy (RT) with an immunocytokine [IC; anti-GD2 monoclonal antibody (mAb) linked to IL-2] cures approximately 50% of B78-D14 tumor-bearing mice, inducing durable, T cell-dependent, tumor-specific immune memory.^25^ Notably, B78-D14s lack MHC-I expression, even after IFN-γ stimulation, but can express MHC-II following IFN-γ treatment.^28,29^ We therefore examined the mechanisms underlying curative immune responses in this MHC-II^+^ B78-D14 tumor model to identify TME factors that support adaptive immunity and that mediate tumor killing. Our studies revealed a critical role for CD4s in response to the RT+IC regimen. We observed that CD4s, including Tregs, engaged in tumor cell trogocytosis, which was accompanied by degranulation, indicating cytotoxic activity. These findings demonstrate that, upon IC treatment, Tregs can trogocytose tumor antigens, acquire effector-like properties, and contribute directly to tumor killing in MHC-II⁺ melanoma.

## Results

### CD4 T cells are required for tumor control

We have previously established a radio-immunotherapy regimen in which radiation therapy (RT) is followed by five consecutive doses of immunocytokine (IC) (Fig. 1A), and shown that RT+IC elicits durable, T cell-dependent antitumor responses in the B78-D14 melanoma model.^25^ B78-D14s have aberrant expression of MHC-I and MHC-II, with discordant results reported and we observe that coculture of B78-D14 with IFN-γ for 48hrs induces expression of MHC-II but not MHC-I (**Fig S1A**).^28–30^ Although IFN-γ treatment can upregulate MHC-II on B78-D14 cells, the resulting expression is heterogeneous, with a subset of tumor cells exhibiting little to no detectable MHC-II by flow cytometry (**Fig. S1A**). Tumor cell expression of MHC-II can augment the response to immunotherapy, as MHC-II⁺ tumor cells can function as antigen-presenting cells, stimulating T helper (Th) and T effector (T_eff_) responses to promote adaptive antitumor immunity.^21,31,32^ *Ex vivo* analyses of B78-D14 tumors following RT+IC revealed increased MHC-II expression in tumors from “responding” mice (based on tumor regression) compared to tumors from non-responders (**Fig. 1B-C, Fig. S1B-D**), suggesting that MHC-II expression may contribute to effective antitumor responses. Because MHC-II interacts with CD4s, these findings raise the possibility that CD4-mediated mechanisms underlie part of the therapeutic efficacy in this model. Indeed, immune cell depletion studies confirmed that CD4s are required for the initial antitumor response: mice depleted of CD4s during RT+IC therapy failed to respond, whereas depletion of CD8s or NK cells did not impair therapeutic efficacy (**Fig. 1D, Fig. S2A**). Conversely, in two other tumor models (NXS2 and MC38) that express MHC-I (**Fig S1A**), antitumor efficacy of radio-immunotherapy (RT+IC in NXS2 or RT+IL-2+anti-CTLA4 for MC38) exhibited a distinct dependency on CD8s, but not CD4s (**Fig. S2B-2C**).^33^ These findings suggest that, in contrast to the MHC-II⁺ B78-D14 model, which depends on CD4s, expression of MHC-I in NXS2 and MC38 promotes CD8-driven antitumor responses. Taken together, these findings are consistent with a model in which MHC-II expression favors CD4-driven antitumor responses, whereas MHC-I expression is associated with CD8-driven responses, although additional factors within the TME likely also influence these dynamics.^34^

**Figure 1:**
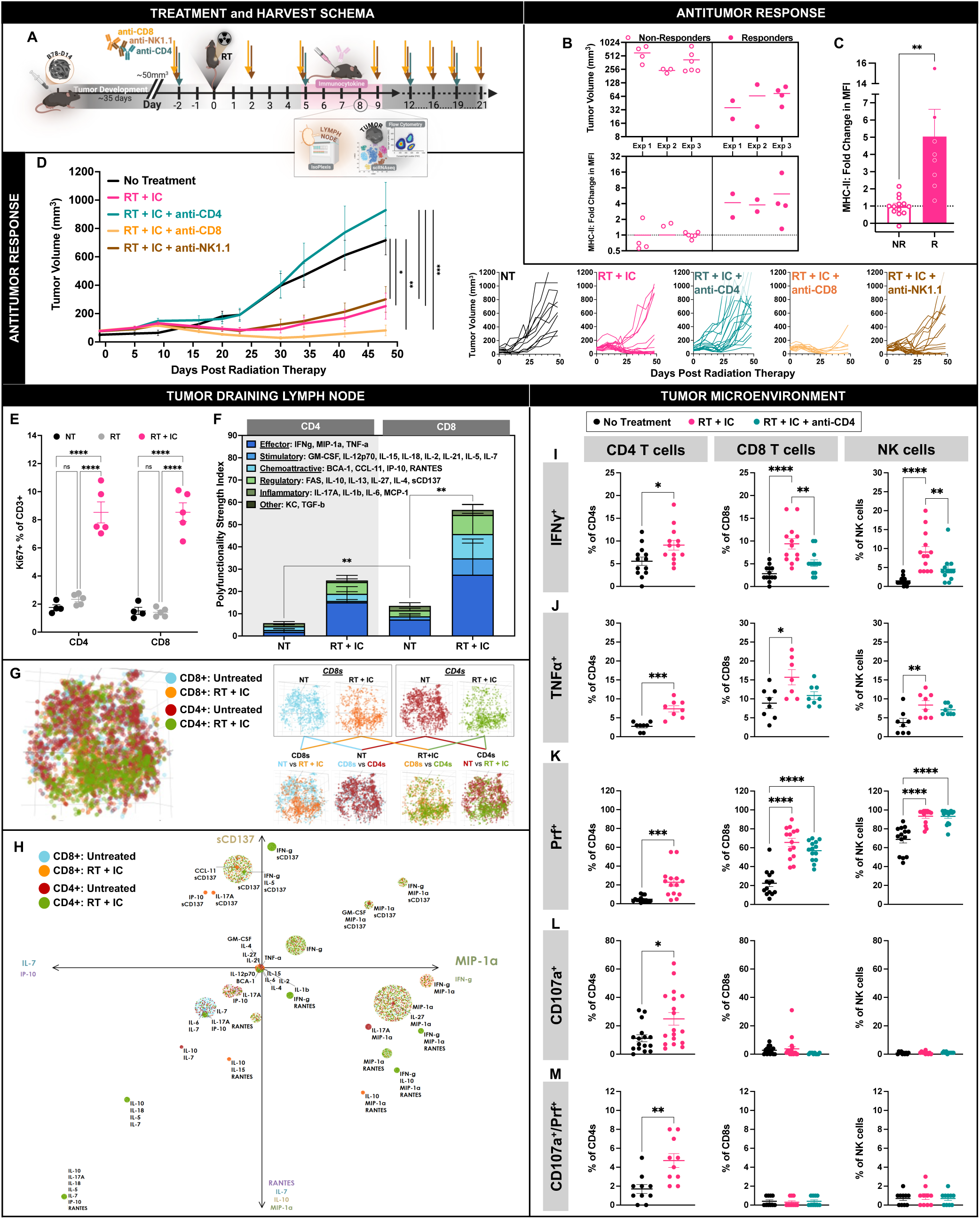
CD4^+^ T cells are Required for Response to RT+IC in B78-D14 Tumors. **(A)** Treatment and harvest schema for mice bearing B78-D14 tumors treated with RT and IC. **(B)** In three independent studies, mice treated with RT+IC were categorized as responders (R) vs. non-responders (NR) based on tumor size at time of harvest (top). Tumors were then analyzed for MHC-II expression (bottom); see additional details in **Fig. S1**. **(C)** Increased tumor MHC-II expression significantly correlated with treatment response. n = 7-11/group; 3 independent experiments combined. **(D)** Immune subset depletion demonstrated that CD4s were required for tumor rejection, while CD8s and NK cells were dispensable. n=11-18/group; 3 independent experiments combined. **(E–H)** From TDLNs harvested on day 7: **(E)** RT+IC increased Ki67 expression in both CD4s and CD8s. **(F)** IsoPlexis analyses showed RT+IC enhanced production of effector cytokines in both T cell subsets, along with increased regulatory cytokines**. (G)** IsoPlexis t-SNE plots showed that RT+IC-CD4s and RT+IC-CD8s clustered together with distinct cytokine profiles compared to NT counterparts. **(H)** Polyfunctional Activation Topology Principal Component Analysis displays cytokine secretion for CD4s and CD8s based on treatment. Each grouping/circle of datapoints represents the same functional group that are randomly offset, but remain within a radius proportional to the secretion frequency of the corresponding group (i.e., large groups = large circles, small groups = small circles). The principal components are labeled according to their correlation with specific cytokines. Here again, both RT+IC-CD8s and RT+IC-CD4s are highly polyfunctional with expression combinations of MIP-1α and IL-27 secreting cells, as well as IFN-γ, CCL-11, IL-5, and sCD137 polyfunctional cells. NT samples pooled from n=3 mice; RT+IC samples pooled from n=3 mice. **(I–J)** From tumors excised on day 8: RT+IC increased IFN-γ (**I**) and TNF-α (**J**) in NK cells, CD4s, and CD8s, but CD4 depletion abrogated this effect in CD8s and NK cells. Prf was increased in all subsets with RT+IC and remained elevated in CD8s and NK cells following CD4 depletion (**K**). However, CD107a expression was restricted to CD4s (**L**), which uniquely co-expressed Prf and CD107a (**M**), indicating direct cytotoxic activity. n = 18/group; 4 independent experiments combined. *p < 0.05, **p < 0.01, ***p < 0.001, ****p < 0.0001; Mean ± SEM.

We next examined the efficacy of T cell priming within the tumor draining lymph node (TDLN) following RT+IC treatment to determine whether CD4s and CD8s were sufficiently activated to attack the tumor. In the TDLN, RT+IC treatment increased proliferation of both CD4s and CD8s (**Fig. 1E**). Single-cell secretory cytokine analyses showed that CD4s and CD8s from RT+IC-treated mice were similarly activated (**Fig. 1F**), with t-SNE plots showing clustering of both treated subsets (CD4s and CD8s) together, distinct from their respective no treatment (NT) counterparts (**Fig. 1G**). This indicates that RT+IC induces a shared cytokine production phenotype in CD4s and CD8s, different from those in NT conditions. Both CD4 and CD8 T cell types from RT+IC groups also exhibited elevated production of effector cytokines compared with NT controls (**Fig. 1F-H**).

To understand the contribution of CD4s to the antitumor response, we depleted CD4s during RT+IC treatment and compared immune activity within the TME to mice that had CD4s. Within the TME, on day 8 after RT+IC treatment initiation, immune cells from RT+IC-treated mice exhibited distinct immunophenotypes with enriched effector subsets which were lost if CD4s were depleted during treatment (**Fig. 1I-M**). This suggests CD4 depletion abrogates RT+IC-induced TME reprogramming. RT+IC significantly increased IFN-γ and TNF-α in CD4s, which could explain the parallel IFN-γ and TNF-α enhancements observed in CD8s and NK cells (**Fig. 1I-J**). However, when CD4s were depleted during RT+IC, IFN-γ and TNF-α production by CD8s and NK cells was significantly reduced, indicating reduced activation. We also examined cytotoxic potential, defined by lytic granule release of perforin (Prf) and CD107a expression. RT+IC increased cytotoxic CD4s in the TME (**Fig. 1K-L**). Although CD8s and NK cells were not required for tumor control against B78-D14s, we observed an increase in Prf-expressing CD8s and NK cells following RT+IC, with NK cells having the highest levels (**Fig. 1K**). Notably, despite increased Prf expression, CD8s and NKs lacked CD107a surface expression, suggesting limited degranulation and cytotoxic activity. In contrast, CD4s were the only subset with increased CD107a⁺ and Prf⁺ cells following RT+IC, implicating cytotoxic CD4s as key effector cells (**Fig. 1M**).

### CD4 T cells mediate both primary and memory responses

In addition to a requirement for CD4s in the initial antitumor response against B78-D14, we interrogated whether CD4s played a role in long-term immune memory. After initial tumor clearance, tumor-free mice were rechallenged with B78-D14 tumors to evaluate immune memory (**Fig 2A**). Mice depleted of CD4s during the rechallenge were unable to reject B78-D14 tumors, whereas depletion of CD8s or NK cells did not impair rejection (**Fig. 2B-C**). These findings suggest that CD4s not only drive the primary antitumor response but are also essential for mediating rejection of the rechallenged tumor, indicating sustained immune memory.

**Figure 2.**
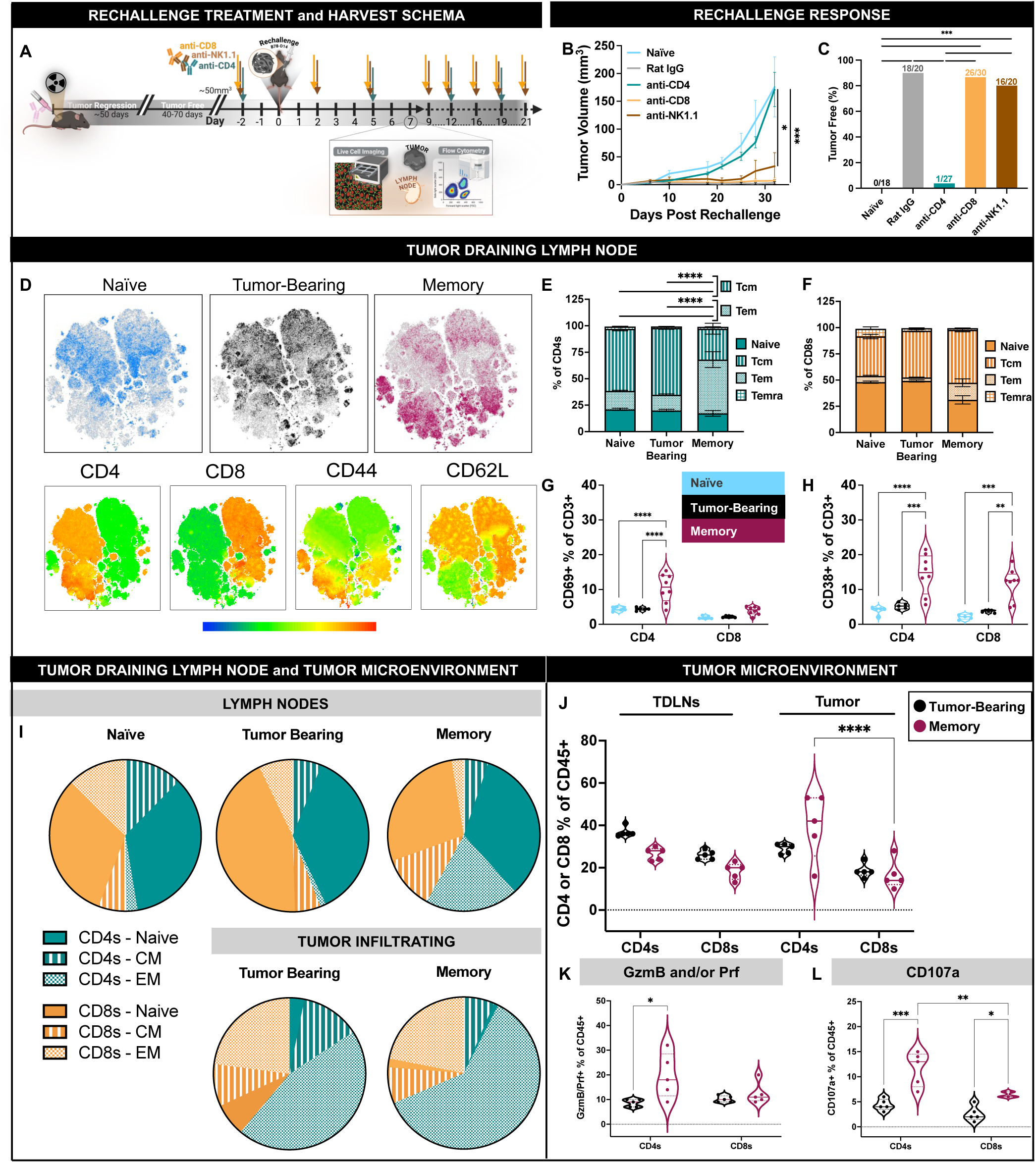
CD4^+^ T cells are Required for the Memory Response in mice cured by RT+IC. **A)** Treatment and harvest schematic for tumor-free mice after RT+IC that were rechallenged with B78-D14s 40-70 days post-clearance. **B-C**) Immune cell depletion during rechallenge revealed that CD4s, but not CD8s or NK cells, were required for rejection, as CD4-depleted mice exhibited tumor growth comparable to naïve controls. n=15-30/group; 2 independent experiments combined. **D)** Flow cytometry of LNs from: age matched naïve mice (i.e., never injected with B78-D14s); or the TDLN 7 days after previously naïve mice were injected with B78-D14s (“tumor-bearing”), or TDLN from memory-mice previously cured of B78-D14 tumors by RT+IC and rechallenged with B78-D14s, with TDLN harvested 7 days after B78-D14 injection, showed shifts in T cell subsets (T_cm_ and T_em_) based on t-SNE analysis of CD4, CD8, CD62L and CD44. **E-H)** Flow cytometry of TDLNs identified memory phenotypes [Naïve, CD44⁻CD62L^+^; T_cm_, CD44⁺CD62L⁺; T_em,_ CD44⁺CD62L⁻; terminally differentiated effector memory cells (Temra), CD44⁻CD62L⁻]. Memory-mice exhibited significantly reduced CD4⁺ T_cm_ and significantly increased CD4⁺ T_em_ (**E**), with no differences found in CD8^+^ memory subsets (**F).** Memory-mice also showed enhanced expression of activation markers—CD69 on CD4s (**G**) and CD38 on both CD4s and CD8s **(H**). **(I)** In TDLNs and the TME, CD4⁺ T_em_ were enriched in memory-mice compared to naïve or tumor-bearing controls. (**J**) CD4s also constituted a greater fraction of the CD45^+^ cells than the CD8s in the TME (but not TDLNs) in memory versus tumor-bearing mice. **(K-L)** Within the TME of memory-mice, CD4s expressed higher levels of GzmB, Prf, and CD107a, consistent with cytotoxic function, whereas CD8s did not. n=5/group; 2 independent experiments. *p < 0.05, **p < 0.01, ***p < 0.001, ****p < 0.0001; Mean ± SEM.

To further characterize T cell responses in the memory setting, we analyzed TDLNs from mice cured of B78-D14 tumors with RT+IC 7 days following rechallenge with B78-D14 (“memory-mice”). Flow cytometry t-SNE analyses showed marked phenotypic changes in both CD4s and CD8s compared with naïve or tumor-bearing (NT) controls (**Fig. 2D**). Upon rechallenge, memory-mice displayed decreased CD4 T central memory (T_cm_) but increased CD4 T effector memory (T_em_) cells (**Fig. 2E**), with no major shifts in the CD8 memory subsets (**Fig. 2F**). Both CD4s and CD8s expressed elevated CD38 during rechallenge, but only CD4s showed significantly increased CD69, suggesting preferential CD4 activation (**Fig. 2G-2H**).^35,36^

We examined LNs from memory-mice in comparison to tumor-bearing (NT) control or naïve mice. In the LNs of naïve mice and the TDLN of tumor-bearing mice, the T_em_ phenotype was more prominent among CD8s than CD4s. In contrast, in the TDLNs of memory-mice, CD4s exhibited a more pronounced T_em_ phenotype compared to CD8s (**Fig. 2I**). Within tumors, memory-mice likewise showed a higher proportion of CD4 T_em_ relative to tumor-bearing controls (**Fig. 2I**). Notably, among CD45^+^ cells in the memory-mice, CD4 and CD8 subset distributions were comparable in TDLNs, whereas the tumor compartment was dominated by CD4s (**Fig. 2J**). Functionally, only CD4s—not CD8s—contained significantly increased cytotoxic granule content (GzmB and/or Prf; **Fig. 2K**) and degranulation (CD107a; **Fig. 2L**) in memory-mice vs. tumor-bearing mice. Furthermore, CD4s from memory-mice had increase CD107a expression compared to CD8s from the same mice (**Fig. 2L**). Together, these results indicate that CD4s are the primary mediators of the memory response to B78-D14 tumors, characterized by a dominant effector memory and cytotoxic phenotype driving long-term protection.

### scRNA-seq identifies phenotypic shifts with functional adaptation of CD4 subsets

To define how RT+IC reshapes CD4⁺ T cell states, we performed scRNA-seq on tumors from NT and RT+IC-treated mice (**Fig. 1A**). RT+IC increased NK and T cell abundance and substantially reduced tumor cells, consistent with treatment-induced tumor clearance (**Fig. 3A**). Within CD4s, we identified six transcriptionally distinct clusters: naïve, T effector cells (T_eff_), T effector memory cells (T_em_), pTregs (proliferative Tregs), eTregs (effector *GzmB-high* Tregs), and oTregs (other Tregs) (**Fig. 3B)** based on their respective gene expression patterns (**Fig. 3C**). Of these six clusters, oTregs were decreased following RT+IC treatment, whereas both T_em_ and eTregs were increased (**Fig. 3D**).

**Figure 3.**
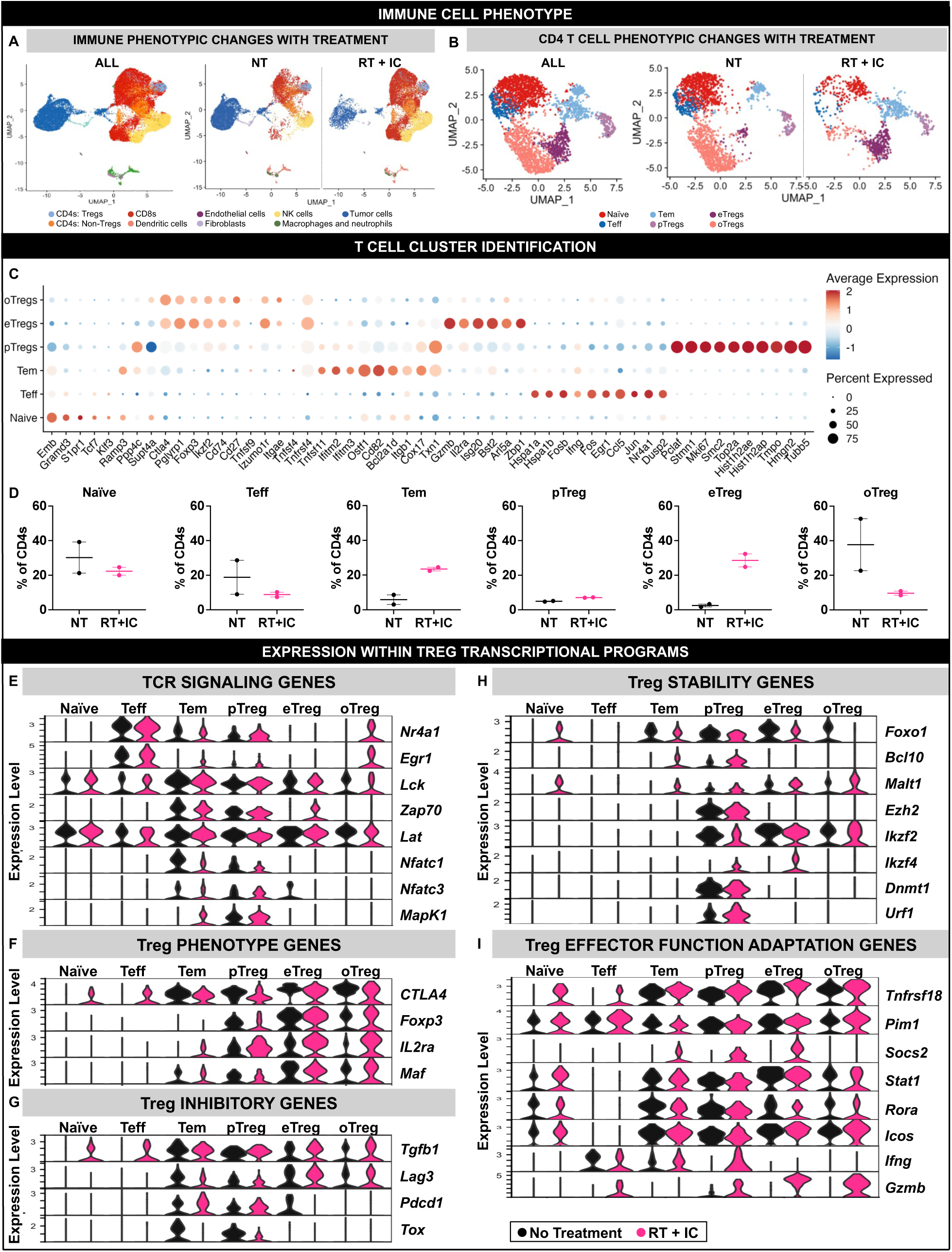
scRNA-seq Reveals CD4 Treg Cells are Engaged in the Antitumor Response. Mice bearing B78-D14 tumors, either untreated (NT) or treated with RT+IC, were assessed by scRNA-seq on day 8. **A)** RT+IC treatment reduced tumor cell abundance and reshaped the immune landscape, with redistribution of CD4⁺ subsets (non-Tregs and Tregs) and increased infiltration of NK cells and CD8s within the TME. Myeloid and dendritic cells remained minimal in both NT and RT+IC tumors. **B)** Focusing on CD4s, six distinct populations were identified: naïve, T_eff_, T_em_, eTregs, oTregs and pTregs. **C)** Subset-specific RNA expression patterns revealed distinct transcriptional profiles for each CD4 population. **D)** With RT+IC, the frequency of T_em_ and eTregs increased, while naïve, T_eff_, and oTregs decreased. (**E-H**) Key gene expression levels were examined in pathways that contribute to: (**E**) TCR signaling; (**F**) Treg phenotype; (**G**) Treg inhibition; (**H**) Treg stability; and (**I**) Treg effector function in Treg and other CD4 subsets following RT+IC. n=2/group; 1 independent experiment.

Given that CD4s are required for both primary B78-D14 tumor control and memory rechallenge response (**Fig. 1C** and **Fig. 2B**), and the association between tumor MHC-II expression and antitumor response (**Fig. 1B**), we first focused on transcriptional programs relevant to CD4 T cell receptor (TCR) signaling (**Fig. 3E**). T_eff_ cells (from both NT and RT+IC mice) expressed high levels of TCR activation genes (*Nr4a1*^37^ and *Erg1*^38^), while T_em_ cells (from both NT and RT+IC mice) expressed proximal signaling of *Lck* and *Lat*, critical for TCR signaling^39^ (**Fig. 3E**). pTregs also showed high expression levels of these genes after RT+IC, including *Nr4a1* and *Egr1,* which are linked to Treg differentiation.^40,41^ Core Treg identity genes, *Ctla4* and *Foxp3*, were decreased with RT+IC treatment in pTregs, while *IL-2Ra* (IL-2 receptor-*a*) was increased across all Treg subsets with RT+IC (**Fig. 3F**). This increased *IL-2Ra* (IL-2R⍺/CD25) expression in the RT+IC group may be due to elevated IL-2 availability in the TME following IC treatment, acting on IL-2Ra-expressing Tregs.^42^ Despite increased *IL-2Ra* expression with RT+IC, none of the Treg subsets showed increased expression of classical inhibitory or suppressive pathway genes such as *Lag3*^43^, *Pcdc1*^44^, *Tgfb1*^45^ with treatment (**Fig. 3G**).

We then evaluated transcriptional programs related to Treg stability and effector adaptation (**Fig. 3H-I)**.^46^ Helios (*Ikzf2*), a key regulator of Treg stability^47^, was expressed in all Treg subsets, but was decreased in pTregs following RT+IC (**Fig. 3H**). Conversely, Eos (*Ikzf4*)^48^, a major regulator of Treg stability, was minimally expressed in each Treg subset. *Foxo1*^49,50^, required for Treg development and function, was decreased with RT+IC in all three Treg populations (**Fig. 3H**). Together, these features suggest that RT+IC promotes Tregs plasticity^51^, and may promote Tregs to adopt effector-like characteristics upon RT+IC. Supporting this, *Socs2,* an essential regulator of Treg plasticity^52^, was increased in all three Treg clusters with RT+IC (**Fig. 3I)**. Genes linked to effector Treg adaptation, including GITR (*Tnfrsf18*)^53^, were upregulated in all three Treg clusters with treatment, with increased *IFN-γ* expression in pTregs and substantially elevated *Gzmb* in all Treg subsets, particularly eTregs (**Fig. 3I**). These features suggest that RT+IC promotes a shift toward effector-like, cytolytic Treg states within the TME.

### Tumor trogocytosis by CD4 T cells correlates with effector function

Following RT+IC treatment, a subset of CD45^+^ immune cells became GD2^+^ **(Fig. 4A)**. GD2, the target antigen of the anti-GD2 IC component of this therapy, is a disialoganglioside that is not often expressed on immune subsets.^54^ Among the CD45^+^ cells, only CD4s, but not CD8s or NK cells, were GD2^+^ (**Fig. 4B**). There was no difference in the amount of tumor-infiltrating CD4s based on treatment, but with RT+IC there was a significant increase in the amount of GD2⁺ CD4s compared to NT (**Fig. 4C**). RT+IC increased CD107a expression on CD4s, with GD2^+^ CD4s expressing significantly more CD107a than GD2⁻ CD4s (**Fig. 4D**). Various methods were applied to validate this observed GD2 expression on T cells as opposed to the presence of doublets (CD4s in direct contact with GD2^+^ tumor cells) or GD2^+^ tumor cells aberrantly expressing CD4. Image Stream flow cytometry revealed that while some CD4s were detected in direct contact with GD2⁺ tumor cells as doublets (**Fig. 4E**), the majority of CD4^+^/GD2^+^ signals were instead isolated CD4^+^ cells that displayed punctate GD2 staining on their membranes (**Fig. 4E**, **Fig. S3A**). *In vitro* coculture of T cells isolated from the LN of a naïve mouse with B78-D14s, followed by sorting and cytospin of GD2⁺CD4⁺ cells, further confirmed that the double-positive population represented CD4s that had acquired GD2, rather than doublets of CD4s bound to GD2^+^ tumor cells or tumor cells that aberrantly expressed CD4 (**Fig. S3B-C**). These data indicated that CD4s acquired GD2 expression via trogocytosis of GD2^+^ tumor membranes.

**Figure 4.**
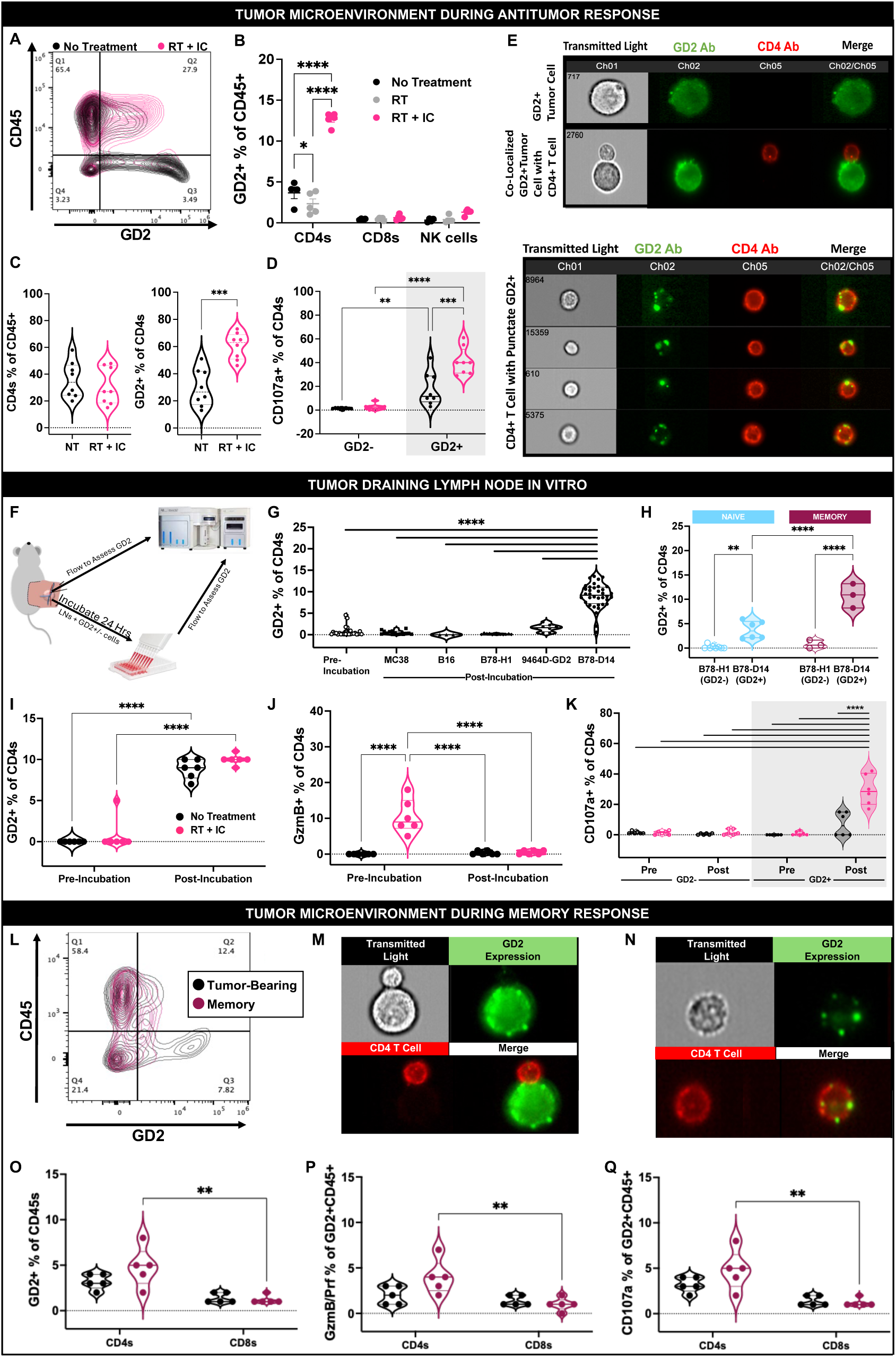
Trogocytosis of B78-D14 by CD4s Influences Effector Cell Function. **A)** Within the TME of B78-D14 tumor-bearing mice, a subset of CD45⁺ cells became GD2⁺ by day 8 after RT+IC (Fig. 1A). **B)** Among immune subsets, CD4s—rather than CD8s or NK cells—displayed the highest GD2 expression following RT+IC. **C)** RT+IC did not alter the overall proportion of CD4s within the TME (left) but significantly increased the frequency of GD2⁺ CD4s (right). **D)** RT+IC increased the percent of CD107a⁺GD2⁺ CD4s compared to NT. No CD107a⁺ cells were detected among GD2⁻ CD4s under either treatment condition. **E)** Image Stream analysis of tumors excised on day 8 revealed (top panel) uniform GD2 expression on tumor cells (anti-GD2 mAb, green), while CD4s were smaller and stained homogeneously with anti-CD4 (red). Some CD4^+^/GD2^+^ signals reflect doublets of green tumor cells adhering to red CD4^+^ cells (top panel), while most reflected a subset of isolated CD4s that displayed bright CD4 signal and punctate GD2 staining (bottom panel), consistent with trogocytosis of GD2 from tumor cells. n=8/group; 2 independent experiments combined. **F)** Schema for TDLNs from B78-D14-bearing mice cocultured with MC38 (GD2⁻MHC-I^+^/MHC-II^lo^), B16 (GD2⁻MHC-I^+^/MHC-II^+^), B78-H1 cells (GD2⁻/MHC-I⁻/MHC-II⁻), 9464D-GD2 (GD2**^+^**MHC-II⁻/MHC-I⁻), or B78-D14 (GD2^+^/MHC-I⁻/MHC-II^+^) cells. **G)** Pre-incubation, CD4s expressed minimal GD2. Post-incubation, GD2 acquisition occurred only with the GD2^+^ B78-D14 coculture, not with other lines. **H)** In similar cocultures, CD4s from TDLNs of tumor-free RT+IC-treated memory-mice showed greater GD2 acquisition than CD4s from naïve controls. **I)** Some CD4s acquire GD2 post-incubation with B78-D14s regardless of *in vivo* treatment status. **J)** Prior to coculture, CD4s from RT+IC-treated mice expressed elevated GzmB compared to NT; post-incubation with B78-D14, GzmB levels returned to baseline. **K)** CD107a was absent from GD2⁻ CD4s but selectively upregulated in GD2⁺ CD4s after B78-D14 coculture, with significantly higher GD2⁺CD107a⁺ frequencies in RT+IC-treated mice. n=3-18/group; 2-5 independent experiments combined. **L)** In the TME of rechallenged memory-mice, a subset of CD45⁺ cells became GD2⁺ by day 7 post-rechallenge. **M-N)** Image Stream confirmed some CD4/tumor doublets with GD2⁺ tumor cells interacting with CD4s (**M**), and isolated CD4s displaying punctate GD2 staining **(N**). **O)** Within the TME post-rechallenge, GD2 expression on CD45⁺ cells was predominantly restricted to CD4s, with higher frequencies in memory versus tumor-bearing mice. **P-Q)** Among GD2^+^CD45⁺ cells, CD4s from memory mice expressed significantly more GzmB/Prf (**P**) and CD107a (**Q**) compared to CD8s, highlighting cytotoxic activity unique to CD4s. n=8/group; 2 independent experiments combined. *p < 0.05, **p < 0.01, ***p < 0.001, ****p < 0.0001; Mean ± SEM.

Trogocytosis involves direct membrane transfer from a donor cell (here, GD2⁺ tumor cells) to a recipient cell (CD4⁺ T cells) during close contact.^55–58^ Trogocytosis by CD4s is primarily thought to be mediated via interaction of the CD4 TCR with MHC-II, most commonly observed through CD4 interactions with antigen-presenting cells. Because B78-D14s express MHC-II (upregulated in responding mice, **Fig. 1B–C**), we hypothesized that CD4 trogocytosis was mediated via TCR:MHC-II interactions. To test this, we incubated TDLN cells from RT+IC-treated mice with various tumor lines differing in MHC and GD2 expression (**Fig. 4F**). Significant GD2 acquisition by CD4s was observed only after coculture with MHC-II⁺GD2⁺ B78-D14 cells (**Fig. 4G**), suggesting that tumor co-expression of both GD2 and MHC-II were needed to facilitate or enhance trogocytosis by CD4s in this setting. Similarly, CD4s from naïve and memory-mice became GD2⁺ only after exposure to B78-D14, with significantly higher acquisition in memory CD4s, consistent with TCR recognition facilitating trogocytosis (**Fig. 4H**).

To determine how trogocytosis was associated with CD4 effector function, CD4s were isolated from TDLNs of control (NT) and treated (RT+IC) B78-D14-bearing mice and co-incubated with B78-D14s (**Fig. 4F**). In contrast to tumor-infiltrating CD4s (**Fig 4C**), TDLN-CD4s were GD2⁻ at baseline but became GD2⁺ following coculture (**Fig. 4I**). TDLN-CD4s from RT+IC-treated mice showed high GzmB expression prior to coculture (pre-incubation), which was lost after incubation (**Fig. 4J**), consistent with degranulation by cells that had released their cytotoxic granules. In line with this, the degranulation marker, CD107a, a separate indicator of recent cytotoxic activity, was increased with coculture specifically in GD2⁺ CD4s (**Fig. 4K**). The amount of GD2 expression on CD4s correlated with CD107a levels (**Fig. S4**), with GD2⁺ CD4s from RT+IC-treated TDLNs expressing significantly higher CD107a than those from NT mice after coculture, further supporting the link between GD2 acquisition and enhanced effector function (**Fig. 4K**). Together, the loss of intracellular GzmB, the presence of CD107a, and the acquisition of GD2 post-incubation, indicate that these GD2^+^ CD4s had degranulated upon target (GD2^+^ tumor cell) engagement and acquired GD2 via trogocytosis.

This phenomenon persisted within the TME of rechallenged memory-mice. Mice cured of primary B78-D14 tumors were subsequently rechallenged with a second B78-D14 injection and TME cells were harvested 7 days later for analysis of CD45⁺GD2⁺ cells (**Fig. 4L**). Similar to trends observed in the initial antitumor response, most GD2⁺ cells in the rechallenge TME were CD4s with punctate GD2 staining, consistent with trogocytosis rather than tumor:T cell doublets (**Fig. 4M-N, Fig. S3D**). CD4⁺ cells remained the predominant CD45⁺GD2⁺ population, with more GD2⁺CD4⁺ cells in rechallenge memory-mice than GD2⁺CD8⁺ cells (**Fig. 4O**). Notably, GzmB expression was significantly increased on GD2^+^ CD4s from the TME of rechallenged memory-mice compared to GD2^+^ CD8s (**Fig. 4P**). GD2⁺ CD4s from rechallenged memory mice exhibited significantly higher CD107a expression compared with GD2⁺ CD8s from the same mice, indicating enhanced degranulation by CD4s during the memory response (**Fig. 4Q**). Collectively, these results demonstrate that CD4⁺ T cells undergo tumor-directed trogocytosis during both primary and memory responses. Acquisition of GD2 correlates with cytotoxic degranulation (CD107a), suggesting that trogocytosis is linked to the effector function of CD4s in this tumor model.

### Treg subsets mediate trogocytosis and acquire cytotoxic potential

Trogocytosis, the direct acquisition of membrane proteins from other cells, can influence CD4 T cell activation and function.^56,59,60^ Because CD4s undergoing trogocytosis with RT+IC treatment (**Fig. 4C&E**) correlated with degranulation, and Tregs exhibited an effector phenotype during treatment (**Fig. 3B–D**), we examined how GD2 acquisition related to the cytotoxic phenotype of CD4 Tregs. Within the TME, treatment-dependent shifts were observed in GD2 acquisition, cytotoxic markers (CD107a, GzmB), and Treg markers (Foxp3, CD25) (**Fig. 5A, Fig. S5**). Following RT+IC, GD2⁺Foxp3⁺ cells were detected in the TME, including both CD25⁺ and CD25⁻ subsets (**Fig. 5B**), with a significant increase in GD2^+^/CD25⁻/Foxp3^+^ cells compared to NT tumors (**Fig. 5B**). GzmB expression was absent in CD4 subsets from NT tumors, regardless of CD25/Foxp3/GD2 expression, but was present in multiple populations following RT+IC—including GD2⁻^/+^/CD25⁻/Foxp3⁻ CD4s, GD2⁻/CD25⁻/Foxp3^+^ CD4s and GD2^+^/Foxp3^+^ CD4s (**Fig. 5C and Fig. S5D**). Following RT+IC, with the exception of GD2^+^/CD25^+^/Foxp3⁻ cells, all GD2⁺ CD4 subsets (CD25⁺Foxp3⁺, CD25⁻Foxp3⁺, and CD25⁻Foxp3⁻) displayed significantly increased CD107a expression, indicating degranulation (**Fig. 5D, Fig. S5E**).

**Figure 5.**
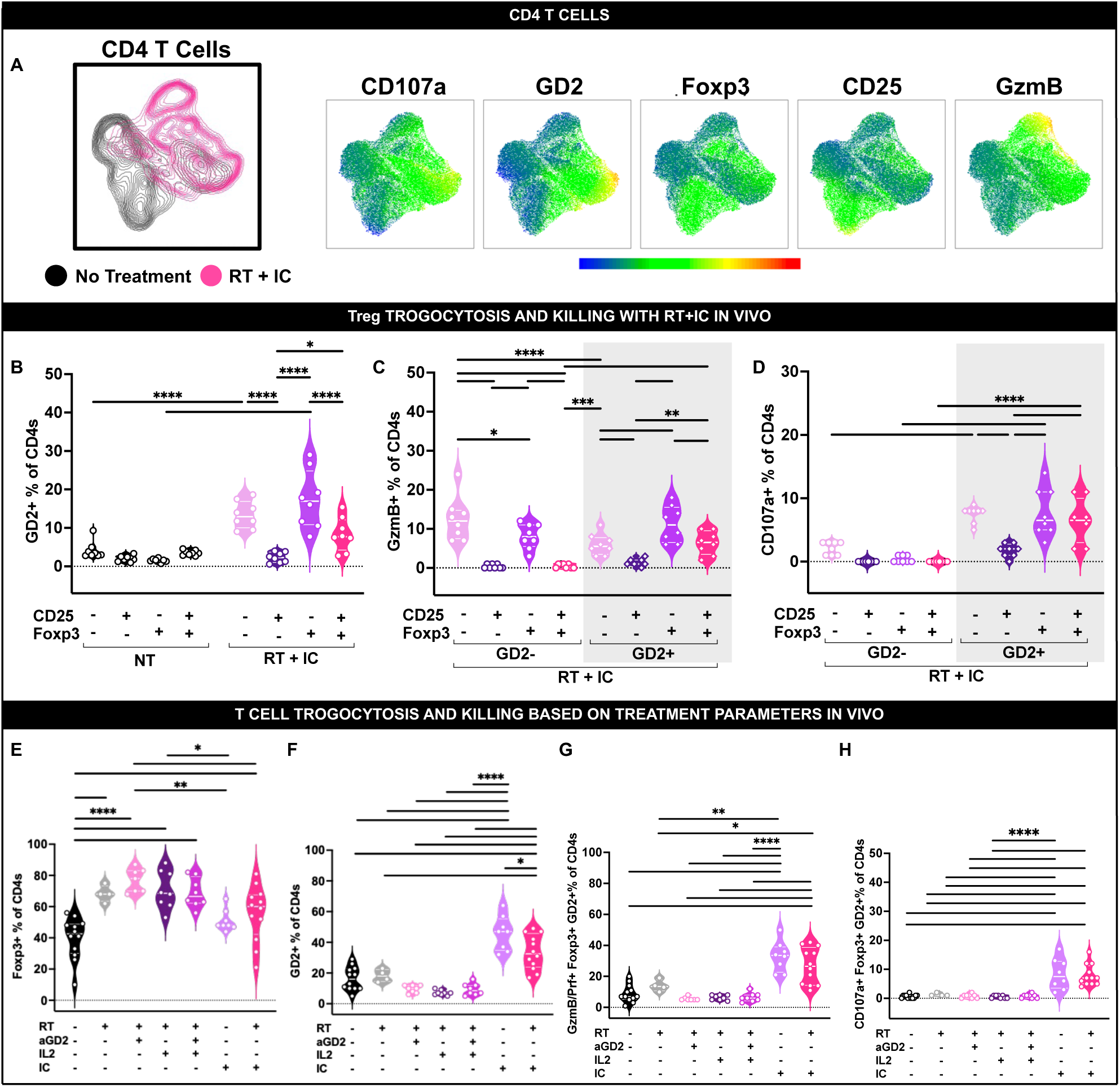
Effector Tregs Trogocytose Tumor Cells and Degranulate. **A)** Flow cytometry of TMEs harvested on day 8 post-treatment revealed treatment-dependent shifts in cytotoxicity markers (GzmB, CD107a) and Treg markers (Foxp3, CD25), aligned with GD2 expression patterns (UMAP analysis, **Fig. S5**). **B)** Following RT+IC, GD2 expression was detected on Foxp3⁺ cells (CD25⁺/⁻) as well as other CD4 subsets. **C)** RT+IC significantly increased GzmB expression across CD4 subsets, including Foxp3⁺CD25⁺/⁻ Tregs. **D)** A subset of Foxp3⁺CD25⁺/⁻ Tregs that became GD2⁺ with RT+IC also showed significantly higher CD107a co-expression compared to GD2⁻ Tregs, with elevated CD107a expression in the RT+IC group restricted to GD2⁺ CD4s (**Fig. S6**). n=8/group; 2 independent experiments combined. **E)** Treatments including RT (RT alone, RT+anti-GD2 (“aGD2”), RT+IL-2, RT+aGD2+IL-2, or RT+IC) increased Foxp3⁺ CD4s compared to NT, whereas IC alone did not. **F)** Only regimens containing IC (IC alone or RT+IC) induced GD2⁺ CD4s in the TME. **G–H)** Similarly, IC-containing treatments (IC alone or RT+IC) drove increased GzmB⁺Foxp3⁺GD2⁺ (**G**) and CD107a⁺Foxp3⁺GD2⁺ (**H**) subsets within CD4s. n=6-12/group; 3 independent experiments combined. *p < 0.05, **p < 0.01, ***p < 0.001, ****p < 0.0001; Mean ± SEM.

To determine which components of the treatment regimen contributed to these phenotypes, we evaluated each element alone or in combination: NT, RT alone, IC (anti-GD2 genetically linked to IL-2), RT+IL-2, RT+anti-GD2mAb, RT+anti-GD2mAb+IL-2, or RT+IC. Treatments including RT significantly increased Foxp3^+^ CD4s compared to NT, whereas IC alone did not (**Fig. 5E**). Only treatments containing IC (IC alone or RT+IC) generated GD2⁺Foxp3⁺ CD4s (**Fig. 5F**), with these groups showing the highest expression of GzmB (**Fig. 5G**) and CD107a (**Fig. 5H**) within the GD2^+^Foxp3⁺ CD4s. Separately, *in vitro* observations (**Fig. 4J**) indicated that some CD4⁺ cells from RT+IC-treated mice expressed GzmB prior to coculture and acquired GD2 during IC-mediated interactions with tumor cells. GD2⁺Foxp3⁺ CD4s isolated directly from the TME exhibited both GD2 and GzmB at harvest, consistent with dynamic regulation of cytotoxic granules during tumor engagement.^61^

### Immunocytokine treatment enables Murine Treg-Mediated Tumor Killing

We then considered whether CD4^+^ T_eff_ cells were being converted to Tregs or whether Tregs were acquiring effector-like properties during RT+IC treatment. To address this, TDLNs from tumor-bearing mice were sorted into Treg cells (CD4^+^CD25^+^CD127⁻) or “NOT Tregs” (CD4^+^CD25^-^CD127^+/^⁻) (**Fig. S6A**).^62^ Analysis of these sorted populations confirmed that the Treg fraction was Foxp3⁺, whereas “NOT Tregs” were Foxp3⁻ (**Fig. S6A**) as CD127 expression inversely correlates with Foxp3.^63,64^ Sorted Tregs were then incubated with B78-D14 tumor cells in media for 24hrs and analyzed by flow cytometry (**Fig. 6A**). Consistent with observations in CD4s (**Fig. 4I)**, CD25⁺Foxp3⁺ CD4s did not express GD2 prior to coculture with B78-D14s (pre-incubation; **Fig. 6B**), but GD2 expression was significantly increased on CD25⁺Foxp3⁺ CD4s post-24hr incubation (**Fig. 6B**). Furthermore, CD107a expression was significantly increased on GD2^+^CD25⁺Foxp3⁺ CD4s compared to GD2⁻CD25⁺Foxp3⁺ CD4s (**Fig. 6C**). Importantly, after coculture, Tregs maintained a CD25⁺Foxp3⁺ phenotype, and “NOT Tregs” did not convert into Tregs (**Fig. S6B**).

**Figure 6.**
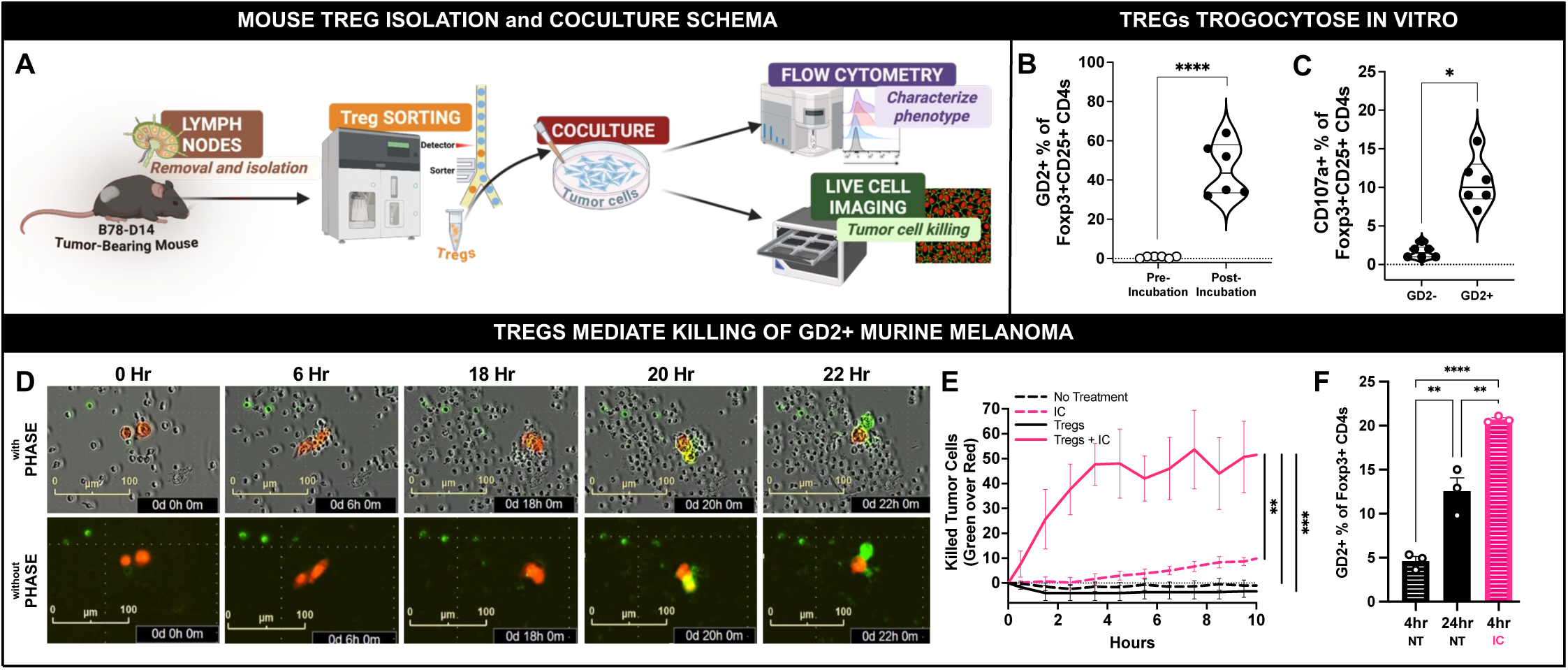
Treg Killing of B78-D14 Tumors is Mediated via IC Treatment. **A)** Schematic of immune cells collected from the TDLN of untreated tumor-bearing mice; Tregs were sorted based on CD4^+^/CD127⁻/CD25^+^ and then incubated with B78-D14+NLS-mKate2 cells in a live cell imaging assay. **B)** Prior to incubation, Foxp3^+^CD25^+^ cells were GD2⁻, but some become GD2^+^ post-incubation with B78-D14s. **C)** The Foxp3^+^CD25^+^ cells that were GD2^+^ had increased CD107a expression as compared to GD2⁻Foxp3^+^CD25^+^ cells after incubation with B78-D14. **D)** Over the course of 22hrs culture with IC, IncuCyte imaging showed that the sorted Tregs (small gray cells) interact with B78-D14+NLS-mKate2 cells (red signal) and cause apoptosis (increased green signal) on the B78-D14s. **E)** Sorted Tregs incubated with B78-D14+NLS-mKate2 cells showed increased killing (as measured by increased green signal of red tumor cells) when co-incubated with IC as compared to Tregs without IC, IC alone (no Tregs), or media alone (no Tregs). n=3 mice/experiment; 2 independent experiments. **F)** Untreated CD4s isolated from spleens of memory-mouse and cocultured with B78-D14s showed a significantly increased frequency of GD2^+^Foxp3^+^ CD4s at 24hrs compared to 4hrs. IC treatment significantly increased GD2 acquisition after 4hrs relative to untreated cells at 4hr or 24hr. n=3 mice/experiment; representative results from 1 of 2 independent experiments. *p < 0.05, **p < 0.01, ***p < 0.001, ****p < 0.0001; Mean ± SEM.

We also monitored the interaction of the sorted Tregs with B78-D14s with or without IC treatment using IncuCyte live-cell imaging (**Fig. 6A**). Over the course of 18hrs following coculture initiation, Tregs trafficked to and surrounded the tumor cells, leading to tumor cell death by 24hrs (**Fig. 6D-E; Video S1-2**). With IC included during treatment, we observed significant induction of B78-D14 killing, with an increased rate of apoptosis of B78-D14s observed within 2hrs post-coculture (**Fig. 6E**). Together with the *in vivo* observation that IC treatment increased expression of GD2 and CD107a on Tregs (**Fig 5B-D**), these results suggest that IC (anti-GD2 linked to IL-2) may promote close interactions between Tregs (via high-affinity IL-2R binding)^65–68^ and GD2⁺ tumor cells (via anti-GD2), enabling both tumor killing and acquisition of tumor-derived GD2 through trogocytosis.

Given our observation of rapid Treg-mediated tumor cell killing (<4hrs) in the presence of IC, and the correlation between GD2 acquisition and cytotoxicity, we next asked whether IC directly enhances GD2 acquisition on the surface of Tregs. While CD4s from memory-mice acquire GD2 following 24hr hours of coculture with B78-D14s (**Fig. 4H**), we tested whether GD2 acquisition could occur over a shorter time frame (4hrs) and whether IC influenced this process. Consistent with our prior findings (**Fig. 4H**), GD2⁺ CD4s were detected after 24hrs of coculture. In contrast, at 4hrs, less than 5% of untreated CD4s became GD2⁺. However, in the presence of IC the extent of GD2 acquisition at 4hrs was significantly greater compared to untreated CD4s at either 4 or 24hrs (**Fig. 6F**).

### Immunocytokine treatment enables human Treg-mediated tumor killing

To determine whether these findings translate to the human setting, T cells isolated from healthy donor human PBMCs were incubated *in vitro* with the GD2⁺ human melanoma cell line, M21—which does express MHC-I and MHC-II with IFN-γ stimulation (**Fig S1A**)—for 4hrs under various treatments (**Fig. 7A**). No differences in the percentages of Foxp3^+^ CD4s were found across treatment groups (**Fig. S8A**). Single-agent treatment with anti-GD2 mAb or IL-2, or the combination of anti-GD2 mAb+IL-2, did not induce GD2 expression on CD4s (**Fig. 7B**), GD2^+^Foxp3^+^ CD4s (**Fig. 7C**), or GD2^+^CD107a^+^ on Tregs (**Fig. 7D**). Conversely, IC treatment led to significant induction in GD2 expression on all CD4s (**Fig. 7B**), including Foxp3^+^ CD4s (**Fig. 7C**), as well as a marked increase in CD107a expression on GD2^+^Foxp3^+^ CD4s (**Fig. 7D**). These effects were abrogated by IL-2R blockade with anti-CD25, confirming that IC directly links Tregs to tumor cells via IL-2R engagement (**Fig. 7B-7D**). To further confirm that these effects occur through direct linkage of Tregs to tumor cells, we evaluated trogocytosis using a GD2/CD3 engager (anti-GD2 × anti-CD3 bispecific antibody, “aGD2×aCD3”). Similar to IC, aGD2×aCD3 treatment resulted in significantly increased expression of GD2 on CD4s and Foxp3^+^ CD4s (**Fig. 7E-F**) and enhanced CD107a expression on GD2^+^Foxp3^+^ CD4s (**Fig. 7G**). These effects were not seen with the antibodies (anti-GD2 mAb or anti-CD3 mAb) given individually and were reduced by anti-CD3 blockade prior to coculture (**Fig. 7E–G, Fig S7B-E**).

**Figure 7.**
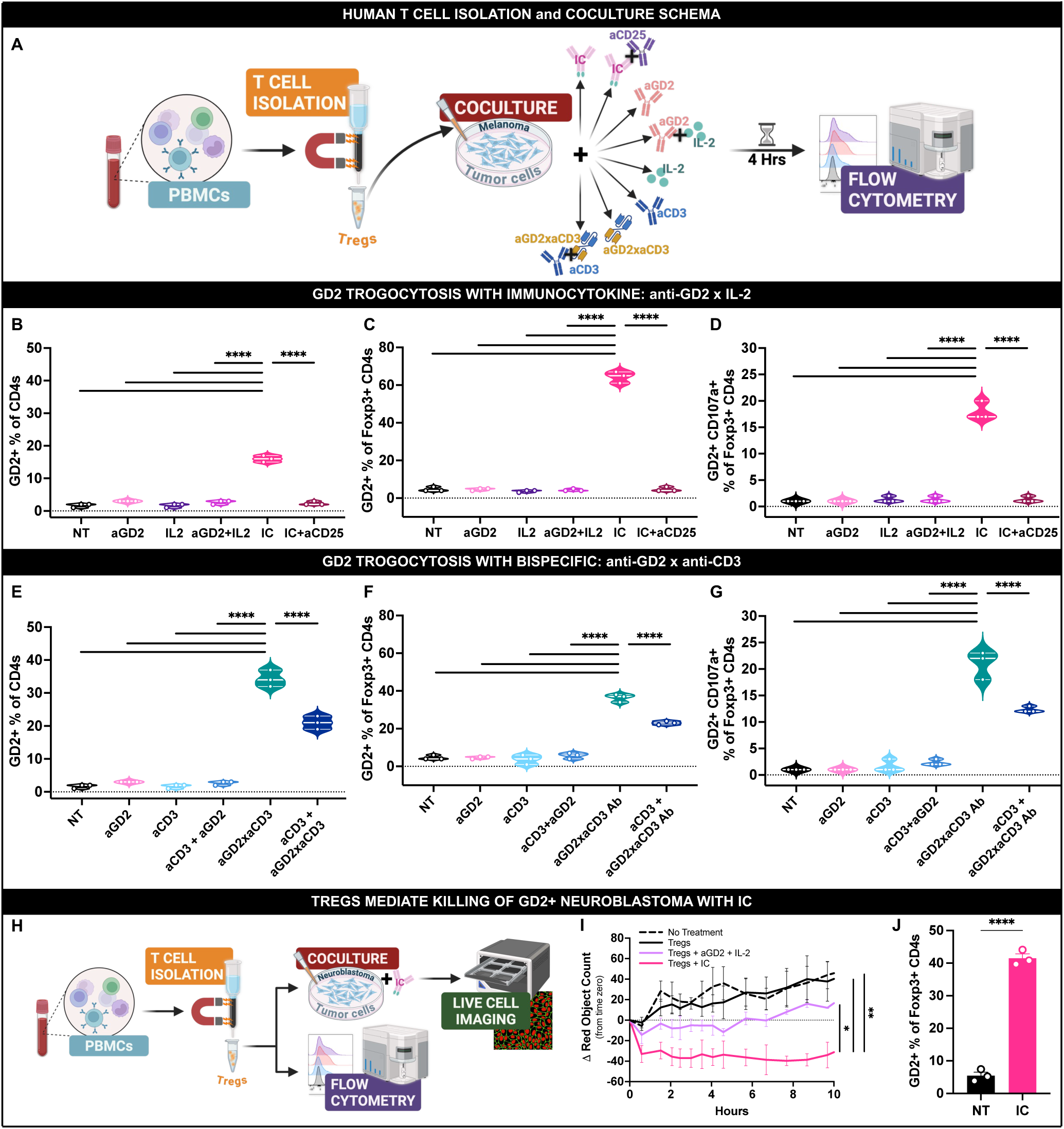
Treg Trogocytosis and Killing of Human Tumors. **A)** Schematic showing T cells isolated from PBMCs cocultured with human GD2^+^ M21 melanoma cells in the presence of aGD2 mAb, IL-2, aGD2 mAb+IL-2, IC, IC+aCD25 mAb (to block the CD25 IL2 receptor), aCD3 mAb+aGD2 mAb, aCD3×aGD2 bispecific Ab, or aCD3×aGD2 bispecific Ab + aCD3 mAb (to block the CD3 receptor) for 4hrs and then analyzed via flow cytometry. **B-C)** Coculture with IC significantly increased the expression of GD2 on CD4s (**B**) as well as on Foxp3^+^ CD4s (**C**). **D)** Coculture with IC also significantly increased CD107a expression on GD2^+^Foxp3^+^ CD4s. aCD25 mAb pretreatment of CD4s, prior to culturing with IC significantly reduced the IC-induced GD2 expression on CD4s and Foxp3^+^ CD4s, as well as decreased CD107a on GD2^+^Foxp3^+^ CD4s (**B,C,D**). **E-F)** Treatment with the aCD3×aGD2 bispecific Ab significantly increased the expression of GD2 on CD4s (**E**) as well as on Foxp3^+^ CD4s (**F**); and aCD3×aGD2 bispecific Ab treatment also significantly increased CD107a expression on GD2^+^Foxp3^+^ CD4s (**G**). These effects were significantly diminished by pretreatment with aCD3 mAb prior to aCD3×aGD2 bispecific Ab treatment, confirming dependence on CD3 engagement (**E-G**). (A compilation of these data can be found in **Fig. S7B-D**). 2 independent experiments. **H)** Schematic showing PBMCs isolated from a healthy donor which were sorted for Tregs by magnetic separation and incubated with human LA-N-1+NLS-mkate2 neuroblastoma cells with or without IC and monitored via live cell imaging. **Fig. S7E** shows efficiency of Treg isolation. **I)** LA-N-1+NLS-mkate2 cells express red fluorescent signal within their nuclei; a change in the signal over time from time zero determines proliferation (increase in cell count) or cell death (decrease in cell count). When LA-N-1+NLS-mkate2 cells are cocultured with Tregs + IC, the LA-N-1 cells show substantial cell death. In contrast, when LA-N-1+NLS-mkate2 cells are cultured alone (No Treatment), or with Tregs alone, or with Tregs together with aGD2 + IL-2, the LA-N-1 cells were proliferating after 8hrs, indicating they were not being killed. **J)** Healthy donor human CD4s cocultured with LA-N-1 tumor cells for 4hrs showed a significant increase in the percentage of GD2^+^Foxp3^+^ CD4s when IC was in the culture compared to untreated (NT). Representative results from 1of 2 independent experiments, 3 separate healthy donors. *p < 0.05, **p < 0.01, ***p < 0.001, ****p < 0.0001; Mean ± SEM.

Finally, to test whether human Tregs are cytotoxic to human GD2⁺ tumors, we cocultured LA-N-1 human neuroblastoma cells—which express MHC-I, but not MHC-II, with IFN-γ stimulation (**Fig S1A**)—with human Tregs in the presence of IC (**Fig. 7H**). Over a 10hr coculture, only Tregs treated with IC were capable of killing LA-N-1 cells (**Fig. 7I**). Because LA-N-1 cells lack MHC-II expression, we next tested whether IC can bypass the requirement for tumor cell MHC-II expression for GD2 acquisition on Tregs. After a 4hr coculture of LA-N-1 cells with healthy donor CD4s treated with or without IC, IC treatment induced GD2 acquisition on Tregs (**Fig. 7J**). Collectively, these results demonstrate that IC coculture with GD2^+^ tumor cells enables mouse and human Foxp3⁺ Tregs to mediate effector-like cytotoxic functions—including GD2 trogocytosis and degranulation—while maintaining their regulatory phenotype, highlighting their capacity for functional adaptation in response to IC-mediated tumor engagement.

## Discussion

External beam RT followed by IC induces curative responses in mice bearing established B78-D14 tumors.^25^ RT not only triggers DNA damage and tumor cell death but also promotes immune activation within the TME. IC amplifies this response by stimulating infiltrating immune cells, including CD4⁺ T cells. In the B78-D14 model—MHC-I–deficient but MHC-II–inducible—CD4s are the primary mediators of RT+IC responses. Direct tumor engagement by CD4s was evident via GD2 trogocytosis, with Foxp3⁺ subsets (Tregs and CD25⁻/Foxp3⁺ cells) comprising ∼1/3 of GD2⁺ CD4s and displaying antitumor effector activity. These findings demonstrate that regulatory populations can participate directly in tumor cytotoxicity.

CD4⁺ T cells are functionally diverse, and their plasticity can be harnessed to drive antitumor responses following immunotherapy.^69–73^ Prior work has demonstrated MHC-II-dependent cytotoxic activity of CD4s against B16 melanoma.^72,74^ Like B16, B78-D14 also express MHC-II under cytokine stimulation (e.g., IFN-γ). Unlike B16, B78-D14 do not express MHC-I, even after IFN-γ. In this context, CD4s take a lead role in the initial antitumor response and are essential in memory (rechallenge) responses. Upon treatment, CD4s exhibited increased activation markers, degranulation molecules, and cytolytic granules. While CD8s and NK cells showed elevated perforin, they did not degranulate, and their activation depended on CD4-mediated signaling, as CD4 depletion abrogated TNF-α and IFN-γ production. Loss of tumor rejection and memory responses following CD4—but not CD8 or NK cell—depletion underscores CD4s as central orchestrators of RT+IC-mediated immunity.

Trogocytosis—the transfer of membrane components and associated antigens between cells—has been described as a means by which T cells dynamically acquire information from antigen-presenting or target cells.^58,75^ GD2 acquisition by CD4s occurs, in part, through an MHC-II-dependent process. CD4s acquired GD2 after *in vitro* coculture only when tumor cell lines expressed MHC-II, and GD2 acquisition was enhanced in memory CD4s. These findings are consistent with a model in which memory CD4s undergo enhanced trogocytosis upon re-encountering MHC-II-presented tumor antigens. This process likely reflects a dynamic mode of antigen engagement that enables CD4s to sustain cytokine production and cytolytic potential.^55,58,60,75^ Beyond MHC-II-dependent processes, IC accelerated GD2 acquisition and cytotoxicity even when target cells lacked MHC-II, indicating an MHC-independent mechanism*. In vitro*, IC accelerated GD2 trogocytosis by CD4s and enhanced tumor killing, even when target cells lacked MHC-II. *In vivo*, during the antitumor response with RT+IC, a subset of Foxp3⁺ Tregs infiltrating the TME acquired GD2, indicating direct contact with GD2^+^ tumor cells. Given that IL-2 preferentially binds to and supports cells expressing the high-affinity IL-2 receptor—including Tregs^55,60^—the IC likely supports Treg persistence and engagement in the TME through IL-2Rα/βγ (or high-affinity IL-2R).^58,75^

The capacity of Tregs to mediate cytotoxicity has been debated. Choi *et al.* used an anti-CD3×anti-EGFRvIII bispecific antibody to report the cytotoxic activity of human Tregs against glioblastoma cells.^76^ However, Koristka *et al*. reported that purified human Tregs did not exhibit detectable cytotoxic activity against tumor cells when engaged with several bispecific antibodies.^77^ Our data clarify this by showing that Tregs demonstrate potent cytotoxic function when bridged to GD2⁺ tumor cells via either anti-GD2×IL-2 IC (mouse and human Tregs) or an anti-CD3×anti-GD2 bispecific engager (human Tregs). Under these conditions, Tregs acquired GD2 through trogocytosis and upregulated CD107a, indicating active degranulation, representing a previously unrecognized mechanism by which Tregs can exert direct tumoricidal activity.^76^ Importantly, these effects were abrogated by IL-2Rα (for IC) or CD3 blockade (for the anti-CD3×anti-GD2 bispecific engager), confirming that direct physical linkage between Tregs and tumor cells via high-affinity IL-2R or CD3 engagement is required. The prominence of Treg engagement in this process may reflect their constitutive expression of the high-affinity IL-2R, which we previously showed is required for IC-mediated bridging of immune cells to tumor cells.^65–68^ In both mice and humans, resting Tregs uniquely express this receptor complex at high levels,^78^ providing a mechanistic explanation for the robust GD2 acquisition, trogocytosis, and cytolytic activity we observed in Tregs relative to other CD4⁺ subsets.

Trogocytosis by Tregs has been described in the context of acquiring the molecules from antigen-presenting cells such as dendritic cells. ^45, 79^ In tumors, Treg cytotoxicity has primarily been associated with suppression of effector lymphocytes,^80^ yet direct tumor cell killing by Tregs has remained largely anecdotal.^76,77^ Here, we identify a previously unrecognized role for Tregs in antitumor immunity: following immune engagement, such as that induced by IC, Tregs can trogocytose tumor antigens and directly kill tumor cells. In our model of MHC-I-deficient, MHC-II-inducible tumor cells, a subset of CD4s mediate trogocytosis *in vivo* during the rechallenge memory response, even without IC, supporting a mechanism in which CD4⁺ T cells directly kill tumor cells that upregulate MHC-II via MHC-II-restricted TCR recognition. Collectively, these findings demonstrate that RT+IC therapy reshapes the tumor immune landscape by engaging CD4⁺ T cells as central coordinators and direct effectors of antitumor immunity. Through cytokine signaling, IL-2-driven activation, trogocytosis, and IC-mediated tumor engagement, Tregs and other CD4 subsets gain cytolytic capacity and contribute to tumor elimination. These results highlight the functional plasticity of CD4⁺ T cells, illustrating how targeted cytokine delivery can repurpose classically suppressive populations toward productive antitumor immunity.^81^

## Methods

### Tumor cells

#### Murine Cell Lines

B78-D14 melanoma cells were obtained from Ralph Reisfeld (Scripps Research Institute), and are derived from B78-H1 cells (obtained from Dr. Matthew Krummel, University of California San Francisco)^82^, which were originally derived from B16 melanoma.^83–85^ The B16F10 (B16) melanoma cell line was originally obtained from Dr. William Ershler (University of Wisconsin-Madison).^86^ MC38 is a mouse colon adenocarcinoma cell line that was obtained from Brad St. Croix (NIH). NXS2 is a mouse neuroblastoma cell line that was obtained from Ralph Reisfeld (Scripps Research Institute). B78 cell lines, B16 cells and MC38 cells were grown in RPMI 1640 (Mediatech) supplemented with 10% heat-inactivated fetal bovine serum (HI-FBS), 2 mmol/L l-glutamine, 100 U/mL penicillin, and 100 µg/mL streptomycin. B78-D14 cells lack melanin, but were transfected with functional GD2/GD3 synthase to express GD2.^83,84^ Periodic treatment with Hygromycin B (50 μg/mL) and G418 (400 μg/mL) was used to maintain GD2 expression of B78-D14 cells. 9464D-GD2 is a GD2-expressing cell line derived from 9464D as previously described.^87^ The parental 9464D cell line is a MYCN-driven neuroblastoma cell line derived from TH-MYCN transgenic mice^88^, and a subclone of the parental 9464D cell line selected for that does not express MHC-I even when stimulated with IFN-γ.^87^ 9464D-GD2 and NXS2 cells were grown in DMEM supplemented with 10% HI-FBS, 2 µM L-glutamine, 1 mM sodium pyruvate, 1X MEM non-essential amino acids, 100 U/mL penicillin, and 100 µg/mL streptomycin. Periodic treatment with puromycin (6 μg/mL) and blasticidin (7.5 μg/mL) was used to maintain GD2 expression of 9464D-GD2 cells.

#### Human Cell Lines

M21 melanoma cells were obtained from Ralph Reisfeld (Scripps Research Institute). LA-N-1 neuroblastoma cells were obtained from Patrick Reynolds (Texas Tech University, Lubbock TX). M21 and LA-N-1 cells were grown in RPMI 1640 supplemented with 10% HI-FBS, 2 mmol/L l-glutamine, 100 U/mL penicillin, and 100 µg/mL streptomycin. Mycoplasma testing via PCR was routinely performed.

### Immune Cell/Tumor Cell Coincubation and Cytotoxicity Evaluation

#### Tumor Draining Lymph Node/B78-D14 Cocultures

B78-D14 melanoma cells transduced to express nuclear-localized mKate2 (B78-D14+NLS-mKate2), were plated at 1,000 cells/well in 20 µL/well in 384-well plates (Corning, #353961) and incubated at 37°C for 24hrs. The following day, lymph nodes (LNs) were harvested from mice that had flank B78-D14 tumors that had not received treatment. Immune cells were isolated from the tumor draining LNs (TDLNs) by pressing the LNs through a 70 μm cell strainer into a 50 mL conical tube using the plunger from a 3 mL syringe. Tregs (CD4⁺CD25⁺CD127⁻) or non-Tregs (CD4⁺CD25⁺/⁻CD127⁺/⁻) were isolated using BD FACSAria II cell sorters. Sorted cells were pelleted, resuspended in RPMI 1640 supplemented with 10% HI-FBS and 2 mmol/L L-glutamine, and plated at 20,000 cells/well in 20 µL together with the B78-D14+NLS-mKate2 cells. To assess cytotoxicity, Annexin V–CF®488A Conjugate (Biotium, #29005R) was added to each well according to the manufacturer’s instructions (final volume 20 µL). Wells were then brought to a total volume of 80 µL by adding either 20 µL of media alone or media containing immunocytokine (IC, 50 ng/mL final concentration). Plates were incubated for 24hrs at 37°C in an IncuCyte SX5 live-cell imaging system.

#### Human Immune Cell/Melanoma Cocultures

LA-N-1 human neuroblastoma transduced to express NLS-mKate2 were plated at 1,000 cells/well in 20 µL/well in 384-well plates (Corning, #353961) and incubated at 37°C for 24 hrs. The following day, peripheral blood mononuclear cells (PBMCs) were isolated from healthy donor whole blood by Ficoll density gradient centrifugation. PBMCs were resuspended in RPMI 1640 supplemented with 10% HI-FBS and 2 mmol/L L-glutamine. Treg cells were isolated using the EasySep Human Treg Cell Isolation kit (Stemcell, #100-1136) per the manufacturer’s protocol and plated at 50,000 cells/well in 20 µL together with the LA-N-1+NLS-mKate2 cells. Wells were then brought to a total volume of 60 µL by adding either 20 µL of media alone or media containing immunocytokine (IC, 50 ng/mL final concentration). Cells were monitored on an IncuCyte SX5 live-cell imager.

### T Cell Coincubation for Flow Cytometry Analysis

#### Human Melanoma Cocultures

For Figure 7A-7G, M21 cells were plated at 10,000 cells/well in 50 µL/well in a 96-well plate and incubated 24hrs at 37°C. After 24hrs, PBMCs were isolated from healthy donor whole blood by Ficoll density gradient centrifugation. PBMCs were resuspended in RPMI 1640 supplemented with 10% HI-FBS and 2 mmol/L L-glutamine. T cells were isolated using the EasySep Human T Cell Isolation kit (Stemcell, #17951) per the manufacturer’s protocol and plated at 50,000 cells/well in 50 µL together with the M21 cells. After 4hrs, PBMCs were collected and assessed by flow cytometry.

For Figure 7J, M21 cells or LA-N-1 cells were plated at 50,000 cells/well in 1 mL/well in a 12-well plate and incubated 24hrs at 37°C. After 24hrs, PBMCs were isolated from healthy donor whole blood by Ficoll density gradient centrifugation. PBMCs were resuspended in RPMI 1640 supplemented with 10% HI-FBS and 2 mmol/L L-glutamine and plated at 500,000 cells/well in 1000 µL together with the M21 cells or LA-N-1, with or without IC at 50 µg/mL. After 4hrs, PBMCs were collected and assessed by flow cytometry.

#### Tumor Draining Lymph Node Cocultures for Flow Cytometry

Tumor cell lines (B78-D14, B78-H1, MC38, B16 or 9464D-GD2) cells were plated in 12-well plates at 200,000 cells/well in 1 mL standard culture media/cell line with 200U of IFN-γ and incubated 24hrs at 37°C. After 24hrs, media was replaced with fresh media without IFN-γ. T cells were isolated as above from LNs of naïve mice, or TDLNs of untreated tumor-bearing mice, tumor-bearing mice treated with radiation therapy (RT)+IC, or rechallenge memory-mice. T cells were suspended in RPMI 1640 supplemented with 10% HI-FBS + 0.05 mM *β-Mercaptoethanol* (final concentration) and co-incubated for 24hrs with tumor cell lines. After 24hrs, T cells were removed from the plate, washed with phosphate buffered saline (PBS), and processed for flow cytometry.

For Figure 6F, B78-D14 cells were plated at 50,000 cells/well in 1 mL/well in a 12-well plate with or without IFN-γ (100 U/ml) and incubated 24hrs at 37°C. After 24hrs, splenocytes were isolated from rechallenge memory-mice by pressing the spleens through a 70 μm cell strainer into a 50 mL conical tube using the plunger from a 3 mL syringe. Splenocytes were resuspended in RPMI 1640 supplemented with 10% HI-FBS and 2 mmol/L L-glutamine and plated at 500,000 cells/well in 1000 µL together with the B78-D14 cells, with or without IC at 50 µg/mL. After 4hrs and 24hrs, splenocytes were collected and assessed by flow cytometry.

### T Cell Coincubation for Cytospin Analysis

B78-D14+NLS-mKate2 cells were plated in 12-well plates at 200,000 cells/well in 1 mL standard culture media/cell line with 100U of IFN-γ and incubated 24hrs at 37°C. After 24hrs, media was replaced with fresh media without IFN-γ. T cells were isolated as above from LNs of naïve mice and suspended in RPMI 1640 supplemented with 10% HI-FBS + 0.05 mM *β-Mercaptoethanol* (final concentration) and co-incubated for 24hrs with B78-D14+NLS-mKate2 cells. Following co-incubation, mkate2⁻/GD2⁺/CD4⁺ T cells were sorted and plated onto glass chamber slides for an additional 24hrs at 37°C. As controls, B78-D14+NLS-mKate2 cells alone and T cells alone (not co-incubated) were also plated on glass chamber slides for 24hrs at 37°C. After incubation, slides were stained with Wright–Giemsa stain and imaged by light microscopy.

### Isoplexis Single-Cell Cytokine Profiling

Single-cell multiplex cytokine profiling was performed using the Isoplexis platform on LNs of naïve mice or B78-D14-tumor bearing mice with no treatment (NT) or RT+IC treatment isolated on day 8 after the start of treatment. Immune cells were isolated from LNs by pressing the LNs through a 70 μm cell strainer into a 50 mL conical tube using the plunger from a 3 mL syringe. Isolated immune cells were washed with PBS and immediately processed for single-cell profiling using the IsoCode Single-Cell Adaptive Immune: Mouse T-cells kit (Isoplexis, ISOCODE-1004-4) per the manufacturer’s protocol. Data were analyzed using IsoSpeak software.

### Animal Tumor Growth Studies

C57BL/6NTac mice (5-7 weeks old) were obtained from Taconic Biosciences. A/J mice (5-7 weeks old) were obtained from Jackson laboratories. After arrival, mice were allowed to acclimate to the facility for 1-2 weeks prior to tumor cell injection.

#### Tumor Implantation and Tumor Growth Monitoring

2×10^6^ B78 cells were injected intradermally into the right flank of C57BL/6NTac mice in a volume of 100 μL PBS.^89^ Tumor growth was monitored via twice weekly caliper measurements. Tumor volume was determined using the calculation: width^2^ x length/2. When tumors reached ∼50-100 mm^3^, mice were enrolled in studies for treatment and randomized into treatment groups. Investigators measuring tumor growth once enrolled in treatment groups were blinded to the treatment mice received. Mice were euthanized when either the length or the width of a tumor reached 20 mm and tracked for survival.

#### Treatment Administration

Each randomized group in any single experiment had equivalent starting tumor volumes. Mice receiving RT were immobilized using custom lead jigs that exposed only the dorsal right flank. External beam RT (single dose of 12Gy) was delivered to the right flank tumor surface on day 0 using an X-RAD 320 cabinet irradiator system (Precision X-Ray). Mice receiving IC (AnYxis Immuno-Oncology GmbH, Vienna, Austria) were injected intratumorally on days 5-9 with 50 μg in 100 μL volume. Mice receiving anti-GD2 mAB were given dinutuximab (Unituxin) at 50 μg/dose with or without IL-2 (Proleukin) at 75,000 U/dose in 100 μL volume.

For immune cell depletion studies, CD8 T cells were depleted via anti-CD8 antibody (clone 2.43, BioXcell, # BE0061) at 50 μg in 500 μL PBS per dose on days −2, 2, 5, 8, 12, 15, 18, 22. CD4 T cells were depleted via anti-CD4 antibody (clone GK1.5, BioXcell, #BE0003-1) at 200 μg in 500 μL PBS per dose on days −2, 5, 12, 18. NK cells were depleted via anti-NK1.1 antibody (clone PK136, BioXcell, # BE0036) 200 μg in 500 μL PBS per dose on days −2, 2, 5, 8, 12, 15, 18, 22. Control mice received normal rat IgG antibody (Sigma-Aldrich) at 200 μg in 500 μL PBS per dose on days −2, 2, 5, 8, 12, 15, 18, 22. For Treg depletion studies, mice were treated with anti-CTLA4 (clone 9D9, BRMS) to deplete Tregs at 200μg in 200 μL PBS per dose. As a positive control, mice were depleted of CD4 T cells using anti-CD4 antibody (clone GK1.5, BioXcell, #BE0003-1) at 200 μg in 200 μL PBS per dose. For these studies depletion antibodies were administered on days −1, 2, 5, 8, 11. Flow cytometry of peripheral blood was performed to confirm antibody depletion was successful.

#### Tumor Rechallenge Studies

After treated mice were cured for 40-70 days, tumor rechallenge studies were performed. For the B78 model, 2×10^6^ B78-D14 cells were injected intradermally into the right abdomen of both memory-mice (previously cured with RT+IC) and age-matched naïve mice. All animal work was performed under an animal protocol approved by the Institutional Animal Care and Use Committee at the University of Wisconsin-Madison. A maximum of 5 mice were housed per cage. Standard chow and water were provided *ad libitum*. Enrichment (nesting material and igloo) was provided to all cages. Male and female mice were used for experiments.

### Single-Cell RNA sequencing Studies

#### scRNA-seq sample preparation and acquisition

C57BL/6NTac mice bearing B78-D14 tumors were left untreated or treated with external beam RT (12 Gy, day 0)+IC (50 μg dose, days 5-8). On day 8, 1hr post-injection of IC, mice were euthanized via CO_2_ and tumors were excised. Two tumors per treatment group were finely chopped and suspended in 2.35 mL of RPMI 1640 + Enzyme D (100 μL), R (50 μL) and A (12.5 μL) (Tumor Dissociation Kit, Miltenyi), transferred to a gentleMACS C Tube (Miltenyi) and placed on a gentleMACS Octo Dissociator with Heaters. On the gentleMACs dissociator, program *m_imp* Tumor_02 was used, after which the tubes were inverted and placed in a shaking incubator at 37°C for 40 minutes, and then returned to the gentleMACS dissociator to run on program *m_imp* Tumor_03. Dissociated tumors were filtered through a 40 μm cell strainer (Corning), washed with 15 mL of RPMI 1640, and centrifuged at 300xg for 10 minutes. After removal of the supernatant, live cells were purified from the tumor cell pellet using a Dead Cell Removal Kit (Miltenyi) according to the manufacturer’s protocol. Cell viability was determined and confirmed to be >90%. The Chromium Next GEM Single Cell 5’ Reagent Kits v2 (Dual Index) protocol (10X Genomics) was followed for scRNA-seq processing. RNA quantity and quality were assessed using High Sensitivity D5000 ScreenTape on an Agilent 4200 TapeStation System (UW Biotech Center). Around 10,000 cells per sample were captured for library preparation, and sequenced on an Illumina NovaSeq6000 (Novogene). Two biological replicates were included in a single experiment.

#### scRNA-seq data analysis

Raw reads were aligned to the mm10 reference genome together with UMI (unique molecular identifier) counting using the Cell Ranger pipeline (v3 10X Genomics). Data were filtered using DoubletFinder^101^ to remove potential doublets. Further filtering included only the cells with low mitochondria contents (≤10%) and more than 200 genes covered by the mapping. To integrate the scRNA-seq, we used a fuzzy clustering-based integration method (Harmony method)^102^ to account for potential technical variance across samples. Downstream analyses were based on Seurat single-cell analysis package^103^ including: principal component analysis with standard deviation saturation elbow plot to select the optimal number of principal components, graph-based clustering based on 20 Principal Components using *FindCluster* with different resolution from 0.1-2 to justify the number of clusters based on representative markers overlaid in the hierarchical tree across different resolution (Clustree R package), differential expression analysis using MAST^104^ implemented in Seurat with the cutoff average log2FC=0.25, adjusted *p*-value cutoff of 0.01, and at least 20% of cell expressed the markers. Differentially expressed genes for each subset (major cell types in Fig 3A and CD4 T cell in Fig 3B) were used to annotate the cell types, including *Sox10* to identify tumor cells, *Cd4* for CD4+ T cells, *Foxp3* for Tregs, *Cd8a* for CD8+ T cells, *Ncr1* for NK cells, *Fn1* for fibroblasts, *Pecam1* for endothelial cells, *Csf3r* for neutrophils, *Csf1r* for macrophages, and *Itgax* as well as *Flt3* for dendritic cells. Visualization with dot plots and violin plots were done using Seurat in R (v4.4.2) and Bioconductor platform.

Single-cell RNA-seq data have been deposited at GEO and are publicly available (GEO code: GSE310495). Any additional information required to reanalyze the data reported in this paper is available from the lead contact upon request.

### Flow Cytometry Sample Preparation and Acquisition

#### Tumor Disaggregation and Flow Cytometry

On day 8 post-treatment start, tumors were collected from mice. Tumors were manually disaggregated in petri dishes by chopping into ∼1 mm pieces using a scalpel. Tumors were transferred to 50 mL conical tubes containing 5 mL of complete RPMI 1640, 2.5 mg of collagenase type IV, 250 μg of DNase, and 1X protein transport inhibitor (PTI, eBioscience). Samples were further disaggregated by placing on a heated shaker at 37°C for 45 minutes. Final single-cell suspensions of tumor samples were obtained by passing through a 70 μm cell strainer. Separately, a spleen was collected from a mouse as a control sample for staining. Immune cells were isolated by pressing the spleen through a 70 μm cell strainer into a 50 mL conical tube using the plunger from a 3 mL syringe. Splenocytes were washed with 10 mL of PBS. For red blood cell lysis, splenocytes were first resuspended in 1 mL of sterile water with gentle vortexing for 10 seconds, after which 49 mL of PBS was added and cells were centrifuged to remove wash. Tumor samples and splenocytes were resuspended in flow buffer (PBS + 2% FBS) with 1X protein transport inhibitor (“PTI”; LifeTechnologies), kept on ice, and protected from light for 4-6hrs. 3×10^6^ cells from each individual tumor sample were added to separate 5 mL flow tubes. Leftover tumor cells were pooled, mixed with splenocytes and used for “fluorescence minus one” control samples. For the live/dead control, 1.5×10^6^ cells were frozen at −20°C, thawed and then mixed with 1.5×10^6^ live cells. Samples were resuspended in PBS+PTI and stained with GhostRed780 Viability Dye (Tonbo Biosciences) for 30 minutes at 4°C, protected from light. Samples were washed with flow buffer+PTI and incubated with Fc block ([TruStain FcX](https://www.biolegend.com/en-us/products/trustain-fcx-anti-mouse-cd16-32-antibody-5683), BioLegend, #101319) for 5 minutes at room temperature to reduce nonspecific binding. Samples were stained with surface antibodies in brilliant stain buffer (BD Biosciences) for 30 minutes at 4°C, protected from light. Samples were washed with flow buffer+PTI and fixed with fixation buffer (eBioscience, **#**00-5523-00) for 30 minutes at 4°C, protected from light. After being washed with 1X permeabilization buffer, samples were stained with intracellular antibodies in permeabilization buffer overnight at 4°C, protected from light. The following day, samples were washed once with permeabilization buffer and once with flow buffer. Samples were acquired on an Attune NxT flow cytometer (Thermo Fisher). Antibody panels can be found in supplemental materials (Table S1).

#### FlowJo Analysis

Data were standardized between timepoints using rainbow beads (Spherotech). Data were analyzed using FlowJo (v10.7.1) and gates were determined using fluorescent minus one controls.

### Statistical Analyses

All details on statistical tests, replicates, etc. can be found within the figure captions.

For statistical analysis of tumor volume, killed tumor cells, and red object count, the time-weighted average (area under the volume-time curve, calculated using trapezoidal method) was calculated for each mouse tumor. Time-weighted averages using ANOVA of the time-weighted average of these values, then pairwise t-tests if the overall test was significant. The time-weighted averages were aggregated using the trapezoidal method. No p-value adjustments were made to adjust for inflated type 1 error rate. Survival curves were compared with pairwise log rank tests and response was evaluated by two-sample tests of proportions. Tumor response rate (i.e., tumor-free vs. tumor-bearing) was compared using a chi-square test then pairwise two-sample tests of proportions if the overall test was significant. Significance was assessed at the α = 0.05 level and no adjustments were made to account for inflated type 1 error rate. Analysis was conducted using R version 4.3.1 (2023-06-16). Differences in primary and rechallenge cures were calculated via Fisher exact test. Mouse analyses were carried out in R v4.1.2.

Flow cytometry results are represented as scatter dot plots showing mean (bar) ± standard error of the mean. Each dot represents a single mouse tumor. A one-way ANOVA was calculated with Šίdák’s multiple comparisons test, α = 0.05 (GraphPad Prism v10.6).

Single-cell RNA sequencing results are represented as UMAPs, heatmaps (relative gene expression), dot plots (average gene expression), and violin plots showing mean ± standard deviation.

### Data Analysis and Schematic Design Software

Data were graphed using GraphPad Prism version 10.6 (GraphPad Software, Inc.). Schematic diagrams of study designs were created in BioRender.com.

## Acknowledgements

This work was supported by the Midwest Athletes Against Childhood Cancer, the Cancer Research Institute, the University of Wisconsin Carbone Cancer Center, the Hyundai Hope on Wheels Foundation, the End Kids Cancer Foundation, Stand Up 2 Cancer, The St. Baldrick’s Foundation, Alex’s Lemonade Stand Foundation, The Band of Parents Foundation, The CURE Childhood Cancer Foundation and by public health service grants R35-CA197078, and P01 CA250972 from the National Cancer Institute. NIH Shared Instrumentation Grants (1S10RR025483-01).

The authors thank the University of Wisconsin Carbone Cancer Center Flow Cytometry Laboratory, supported by P30 CA014520, for use of its facilities and services. We additionally thank Dr. Johannes Schoeneberg for his valuable assistance with video stabilization.

## Inclusion and Ethics Statement

All of the coauthors in this study agree that the research conducted is relevant, and they have fulfilled the criteria for authorship required by Nature Portfolio journals, as their participation was essential for the design and implementation of the study. The roles and responsibilities were agreed among collaborators ahead of the research. All of the research conducted was approved under approved Biological Safety, University of Wisconsin Institutional Review Board and Institutional Care and Animal Use Committee for the University of Wisconsin, Madison. This research was not severely restricted or prohibited in the setting of the researchers, and does not result in stigmatization, incrimination, discrimination or personal risk to participants. Local, regional and international research relevant to our study was taken into account in citations.

## Declaration of generative AI and AI-assisted technologies

No AI or AI assisted technologies were used for any component of this research.

