## Supplemental Figures and Table for "Immune-mediated Engagement of T Regulatory Cells with Tumor Cells Results in Trogocytosis and Tumor Cell Killing"

**Supplemental Figure 1. MHC-II expression on B78-D14 tumors correlates with treatment response.** **A)** Expression of MHC-I and MHC-II following 48hr IFN-γ stimulation on murine tumor cell lines (B78-D14, MC38, NXS2, and 9464D-GD2) and human tumor cell lines (M21 and LA-N-1) used in this study. **B**) Tumor-bearing mice were treated with RT+IC. Tumors were collected for flow cytometry from 3 different experiments at time points in which tumor growth differentiation between mice responding (tumors continued to shrink) vs non-responding (tumors began to grow) was observed (day 29 for Exp 1 & 2, day 53 for Exp 3). **C**) MHC-II expression was assessed on tumor cells by gating on $CD45^{-}$ followed by gating on $GD2^{+}$ for MHC-II expression based on fluorescence minus one gating (GD2 on x-axis; MHC-II on y-axis). **D**) MHC-II expression graphs for tumors from responders and non-responders for each individual experiment.

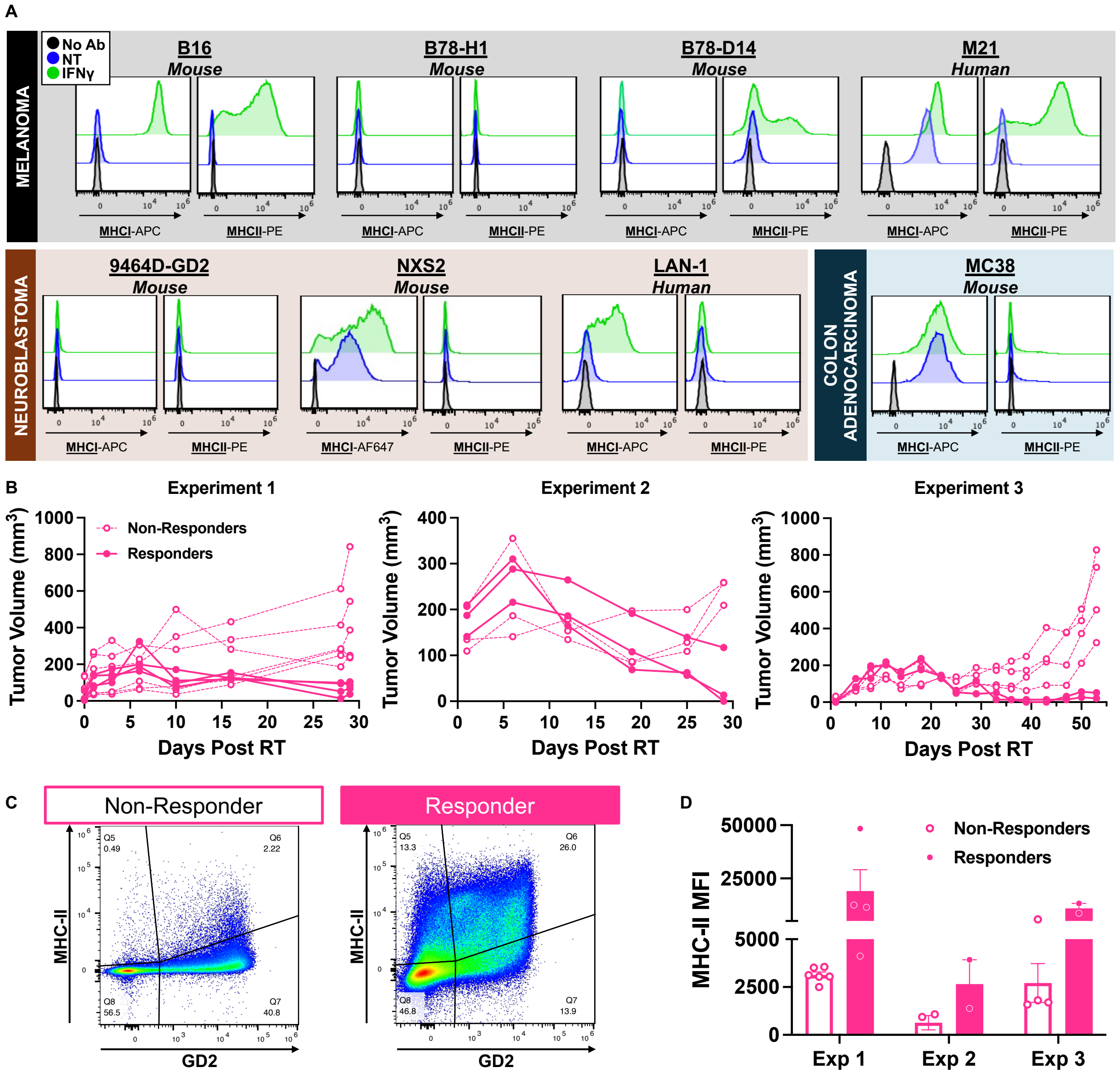

**Supplemental Figure 2. Antitumor response to depletion of CD4 or CD8 T cells depends in part on MHC molecule expression on the tumor cells. A**) Mice with B78 tumors depleted of CD8s or NK cells still respond to treatment with RT+IC, but mice do not respond if depleted of CD4s. **B**) A/J mice bearing NXS2 tumors respond to RT+IC similarly if not depleted or if depleted of CD4s or NK cells; if depleted of CD8s, these mice are less likely to respond. **C**) C57Bl/6 mice bearing MC38 tumors respond to RT+IL-2+anti-CTLA4, but if depleted of CD8s, they do not.[33]

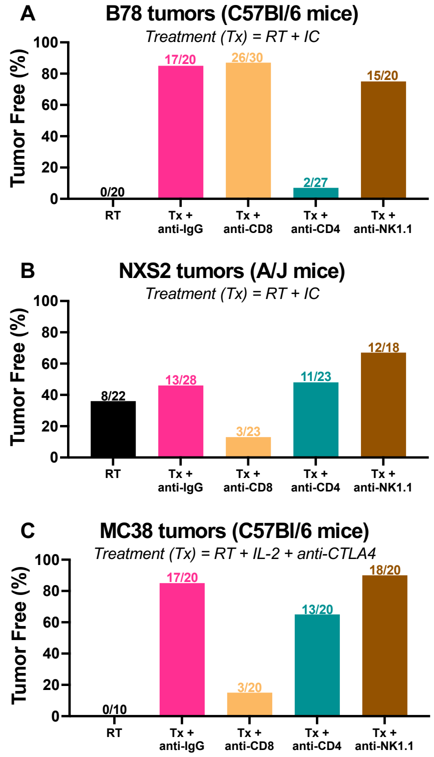

**Supplemental Figure 3. GD2/CD4 co-expressing cells are CD4 T cells that have GD2 expression. A**) Of the GD2$^+$/CD4$^+$ cells identified in the RT+IC-treated TME following treatment with RT+IC (**Fig 4E**), the majority were identified by Image Stream flow cytometry as CD4s containing fragments of GD2 rather than a GD2 cell in direct contact with a CD4 T cell (i.e., a doublet). **B**) Following a 24hr *in vitro* coculture of B78-D14s with T cells isolated from the LN of a naïve mouse, sorted GD2$^+$ CD4s were plated on glass slides. 24hrs later, the slides were stained with Geimsa (top photo) allowing measurement of the GD2$^+$ CD4s, which were 6-7 μm in diameter. Separately, pure B78-D14s cells obtained from culture were plated on glass slides and stained with Geimsa (bottom photo) and found to be 18-21 μm in diameter. **C**) Sorted GD2$^+$ CD4s all appeared to be lymphocytes, with consistent size (~7 μm) smaller than the diameter of isolated pure B78-D14 tumor cells (~19 μm). **D**) Of the GD2$^+$/CD4$^+$ cells identified from the TME during memory rechallenge response, the majority were identified as CD4s with punctate GD2 as compared to doublets of a GD2$^+$ cell in direct contact with a CD4 T cell.

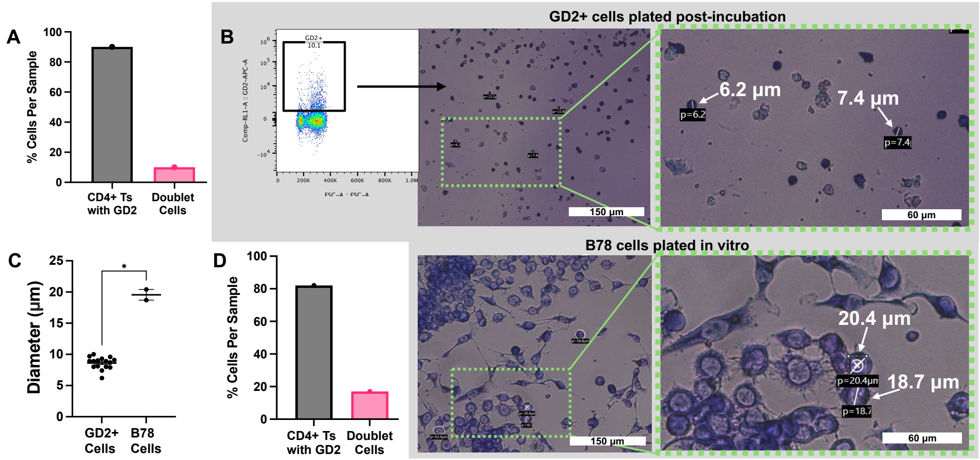


**Supplemental Figure 4. Expression of CD107a on CD4 T cells correlates with the amount of GD2 expression on CD4 T cells.** As shown in Fig 4I&K, TDLN-CD4s from RT+IC-treated or untreated tumor-bearing mice cocultured with B78-D14 cells for 24hrs showed acquisition of GD2 after coculture and became CD107a$^+$. The relationship of GD2 and CD107 expression levels were analyzed. **A**) Increased expression of GD2 has a significant positive correlation with CD107a on CD4s, with RT+IC-treated samples having higher expression of both markers. **B**) Non-expression of GD2 on CD4s has a significant negative correlation with the amount of expression of CD107a on CD4s. Each point represents data from a separate mouse.

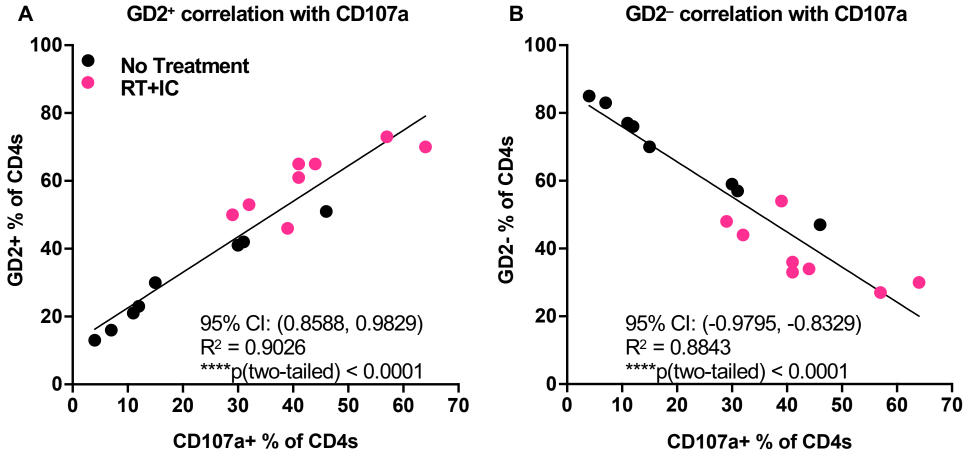

**Supplemental Figure 5. Unbiased FlowSOM Clustering Identifies Treatment-Driven CD4⁺ T-Cell Subsets and Cytotoxic Marker Expression After RT+IC. A**) Unbiased cluster analysis identifies 8 populations based on expression levels of GD2, CD107a, GzmB, Foxp3, and CD25, combining data from untreated (NT) and treated (RT+IC) CD4s in the TME as plotted on UMAP_4LZK using FlowSOM clustering technique. The color intensity in the heat map (right panel) indicates the level of expression of each marker when all samples are combined. **B**) Each individual population is plotted on UMAP_4LZK. **C**) UMAP_4LZK plots show where untreated CD4s (top panel) and RT+IC CD4s (bottom panel) cluster based on expression levels of GD2, CD107a, GzmB, Foxp3 and CD25.**D**) GzmB is expressed on CD25$^{+/-}$/Foxp3/GD2$^{+/-}$ CD4 T cell subsets taken from the TME 7 days after RT+IC treatment. **E**) CD107a is expressed on CD25$^{+/-}$/Foxp3/GD2$^{+/-}$ CD4 T cell subsets taken from the TME 7 days after RT+IC treatment. Two-way ANOVA with Tukey's multiple comparison test. *p < 0.05, **p < 0.01, ***p < 0.001, ****p < 0.0001; Mean ± SEM.

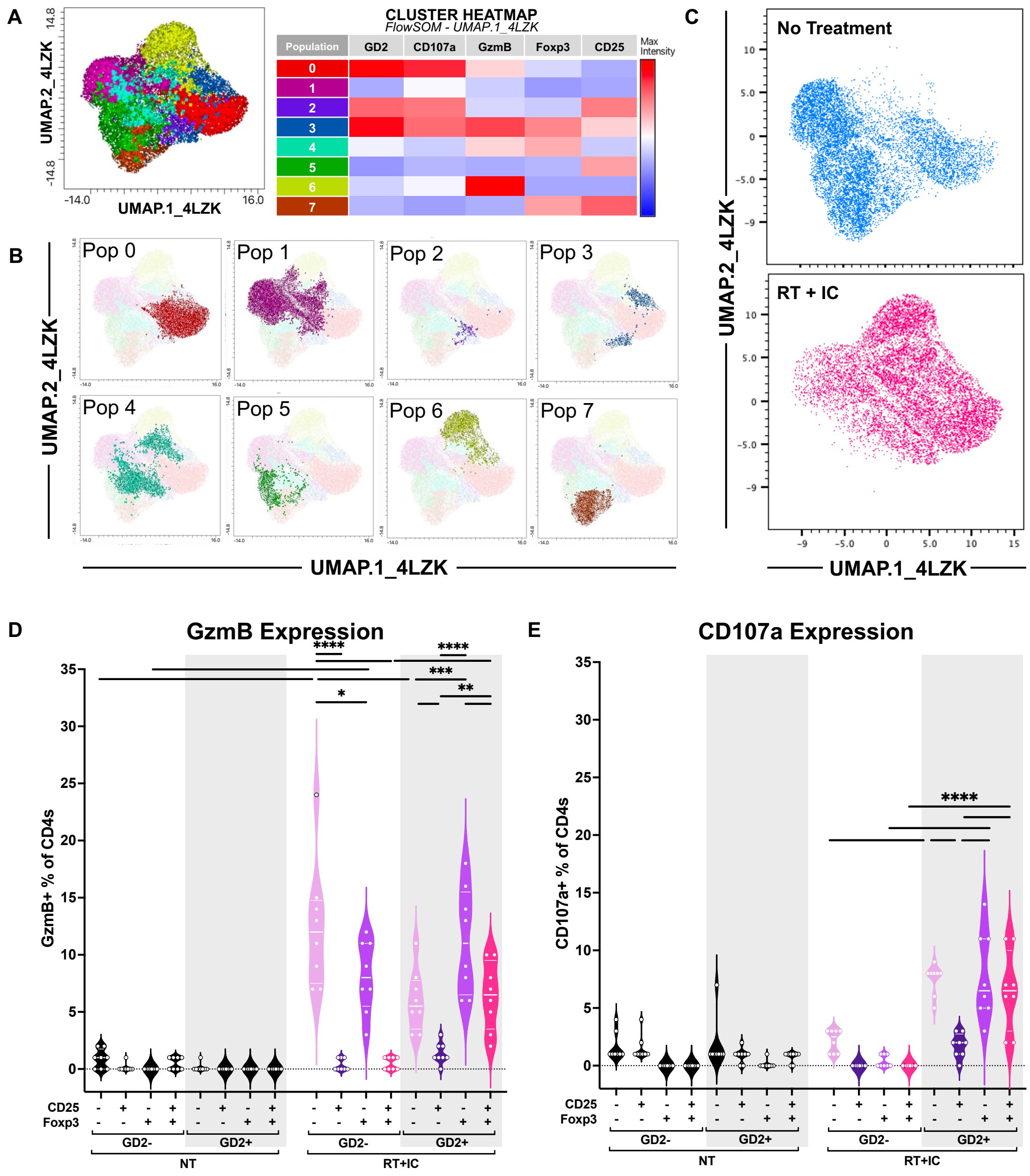

**Supplemental Figure 6. Phenotype of Tregs sorted for the *in vitro* assay. A**) Tregs were sorted from TDLNs of untreated tumor-bearing mice, based on $CD4^{+}CD25^{+}CD127^{-}$. ”NOT” Tregs were sorted based on $CD4^{+}CD25^{+}CD127^{+}$ or $CD4^{+}CD25^{-}$ (left panel). Post-sort (right panel), ~92% of sorted “NOT” Tregs were $CD25^{-}Foxp3^{-}$ with ~5% showing $CD25^{+}Foxp3^{-}$ (top right panel). ~91% of sorted “Tregs” were $CD25^{+}Foxp3^{+}$ (bottom right panel) with ~9% $CD25^{-}Foxp3^{+}$. **B**) Post-coculture with B78-D14 cells, sorted “Tregs” maintain expression of CD25 and Foxp3. “NOT” Tregs remain largely $CD25^{-}Foxp3^{-}$ with a small subset of cells expressing CD25 without co-expression of Foxp3.

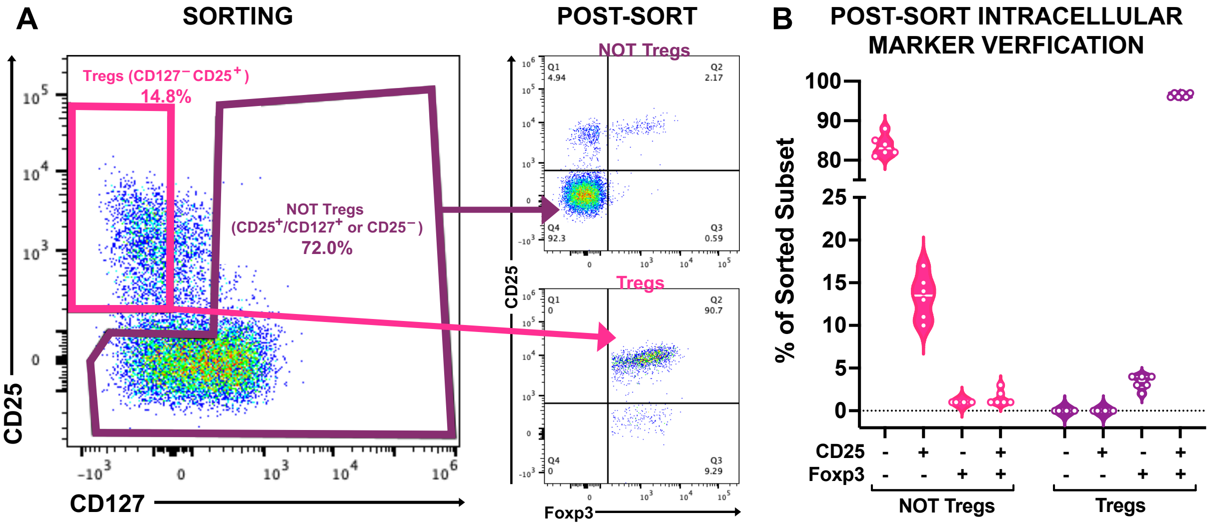

**Supplemental Figure 7. Coculture of human Tregs with tumor cells and IC or aGD2×aCD3 bispecific Ab influences Treg trogocytosis and CD107a expression. A-D**) T cells isolated from human PBMCs cocultured with M21 human melanoma cells were treated with aGD2, IL-2, aGD2+IL-2, IC, IC+aCD25 (to block the CD25 IL-2 receptor), aCD3+aGD2, aGD2×aCD3 bispecific Ab, or aGD2×aCD3 bispecific Ab+aCD3 (to block the CD3 receptor) for 4hrs and analyzed via flow cytometry. **A**) The frequency of Foxp3$^+$ CD4s remained unchanged across treatment conditions. **B**) Only coculture with antitumor antibodies linked to immune cell-targeting domains (IC or aGD2×aCD3) increased the proportion of GD2$^+$ CD4s; this effect of IC was abrogated by aCD25 blockade and for aGD2×aCD3 was partially reduced by aCD3 blockade. The aGD2×aCD3 treatment produced the most GD2$^+$ CD4s. **C**) Similarly, coculture with IC and aGD2×aCD3 increased GD2 expression on Foxp3$^+$ CD4s, with aCD25 abrogating this effect for IC and aCD3 blockade slightly reducing this effect for aGD2xaCD3. IC treatment resulted in the highest GD2 expression on Foxp3$^+$CD4s. **D**) Coculture with IC and aGD2×aCD3 also enhanced CD107a expression on GD2$^+$Foxp3$^+$ CD4s, indicative of degranulation and cytotoxic potential; this IC effect was abrogated by aCD25 and this aGD2xaCD3 effect was modestly reduced by aCD3 blockade. **E)** Efficiency of Treg isolation from human PBMCs by magnetic separation shown in **Fig. 7H&I**.

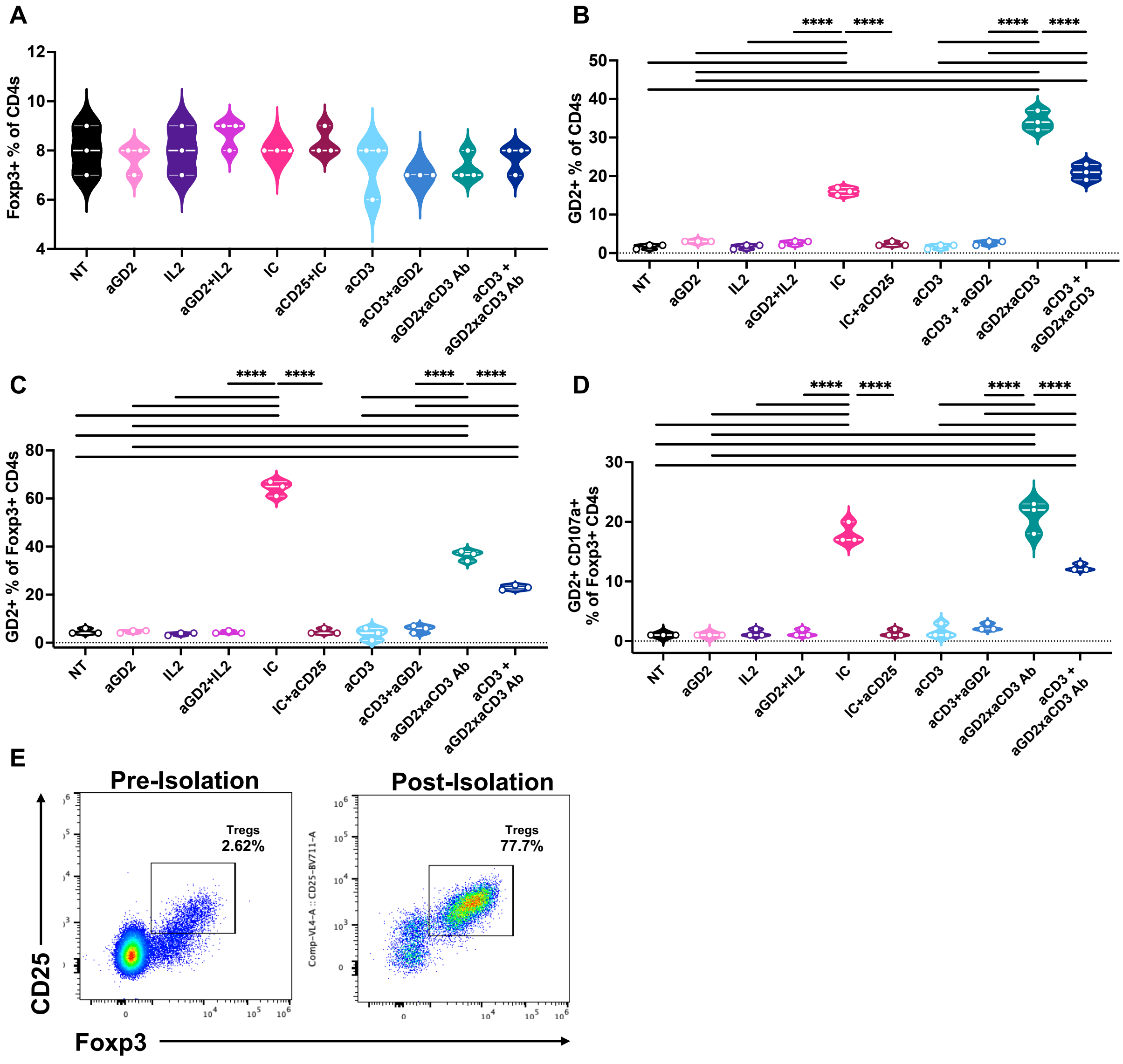

### Supplemental Table 1. Antibodies and live/dead dyes used for flow cytometry studies.

**Mouse Antibodies**

| Target | Fluor | Company | Catalog Number |
|---|---|---|---|
| CD107a | APC | BioLegend | 121614 |
| CD107a | BV421 | BioLegend | 121618 |
| CD127 | BV510 | BioLegend | 135033 |
| CD127 | PE Dazzle | BioLegend | 135032 |
| CD25 | BV711 | BioLegend | 102049 |
| CD3 | PE | BioLegend | 100206 |
| CD3 | PE-Cy5 | BioLegend | 100274 |
| CD38 | PE-Cy7 | BioLegend | 165612 |
| CD4 | AF700 | BioLegend | 100430 |
| CD4 | APC | BioLegend | 100412 |
| CD4 | PE-Cy7 | BioLegend | 100422 |
| CD4 | eFluor450 | LifeTech | 48-0041-82 |
| CD4 | FITC | BioLegend | 100406 |
| CD4 | BV605 | BioLegend | 100451 |
| CD4 | PE | BioLegend | 100408 |
| CD44 | PE-Dazzle | BioLegend | 103056 |
| CD45 | FITC | BioLegend | 103108 |
| CD45 | AF700 | BioLegend | 103128 |
| CD62L | BV711 | BioLegend | 104445 |
| CD69 | BV605 | BioLegend | 104530 |
| CD8 | BV605 | BioLegend | 100744 |
| CD8 | APCR700 | BD Biosciences | 565192 |
| CD8 | FITC | BioLegend | 100706 |
| CD8 | BV711 | BioLegend | 100748 |
| CD8 | APC | BioLegend | 100712 |
| Foxp3 | PECy7 | LifeTech | 25-5773-82 |
| Gzmb | AF647 | BioLegend | 372220 |
| Gzmb | BV510 | BioLegend | 396432 |
| Gzmb | APC | BioLegend | 372204 |
| IFNg | PE-Cy7 | BioLegend | 505826 |
| NK1.1 | PE-Dazzle | BioLegend | 108748 |
| NK1.1 | PE-CF594 | BD Biosciences | 562864 |
| Perforin | PE | BioLegend | 154306 |
| TNFa | BV421 | BioLegend | 506328 |
| TNFa | BB700 | BioLegend | 566510 |
| MHC-I (H2Kb/Db) | AF647 | BioLegend | 114612 |
| MHC-II (I-A/I-E) | PE | BioLegend | 107608 |
| MHC-I (H-2Dd) | APC | LifeTech | 17599980 |

**Human Antibodies**

| Target | Fluor | Company | Catalog Number |
|---|---|---|---|
| GZMB | AF647 | BioLegend | 515406 |
| GD2 | APC | BioLegend | 357306 |
| CD4 | APC | BioLegend | 344614 |
| CD4 | APC R700 | BD Biosciences | 566808 |
| GD2 | APC-Fire750 | BioLegend | 357322 |
| GD2 | BB700 | BD Biosciences | 745751 |
| CD107a | BV421 | BioLegend | 328626 |
| CD8 | BV605 | BioLegend | 344742 |
| GD2 | FITC | BioLegend | 357314 |
| GD2 | PE | BioLegend | 357304 |
| Foxp3 | PE | BioLegend | 12-4776-42 |
| GD2 | PE-Cy7 | BioLegend | 357308 |
| Foxp3 | PE-Cy7 | LifeTech | 25-4776-42 |
| MHC-I (HLA-A,B,C) | AF647 | BioLegend | 311414 |
| MHC-II (HLA-DR, DQ, DP) | PE | BioLegend | 361716 |

**Live Dead Dyes**

| Target | Fluor | Company | Catalog Number |
|---|---|---|---|
| Live/Dead | Ghost Dye 510 | Cytek | 13-0870-T100 |
| Live/Dead | Ghost Dye 780 | Cytek | 13-0865-T100 |
| Live/Dead | Ghost Dye 516 | Cytek | 13-0867-T100 |
| Live/Dead | Dapi | LifeTech | D21490 |
