## Supplemental Videos for "Immune-mediated Engagement of T Regulatory Cells with Tumor Cells Results in Trogocytosis and Tumor Cell Killing"

### Slide 1
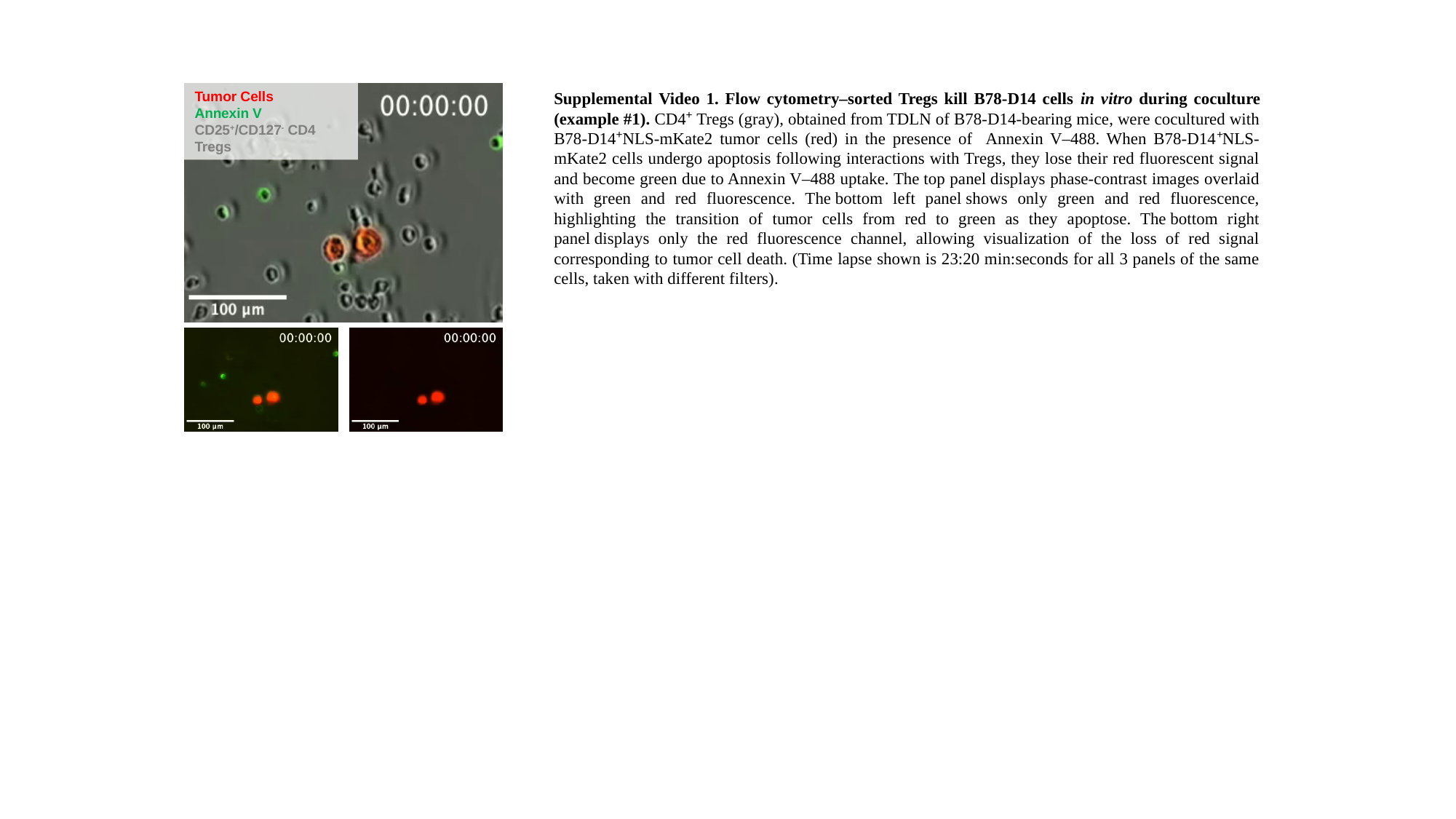

Tumor Cells
Annexin V
CD25^+^/CD127^-^ CD4 Tregs
Supplemental Video 1. Flow cytometry–sorted Tregs kill B78-D14 cells in vitro during coculture (example #1). CD4⁺ Tregs (gray), obtained from TDLN of B78-D14-bearing mice, were cocultured with B78-D14⁺NLS-mKate2 tumor cells (red) in the presence of Annexin V–488. When B78-D14⁺NLS-mKate2 cells undergo apoptosis following interactions with Tregs, they lose their red fluorescent signal and become green due to Annexin V–488 uptake. The top panel displays phase-contrast images overlaid with green and red fluorescence. The bottom left panel shows only green and red fluorescence, highlighting the transition of tumor cells from red to green as they apoptose. The bottom right panel displays only the red fluorescence channel, allowing visualization of the loss of red signal corresponding to tumor cell death. (Time lapse shown is 23:20 min:seconds for all 3 panels of the same cells, taken with different filters).

### Slide 2
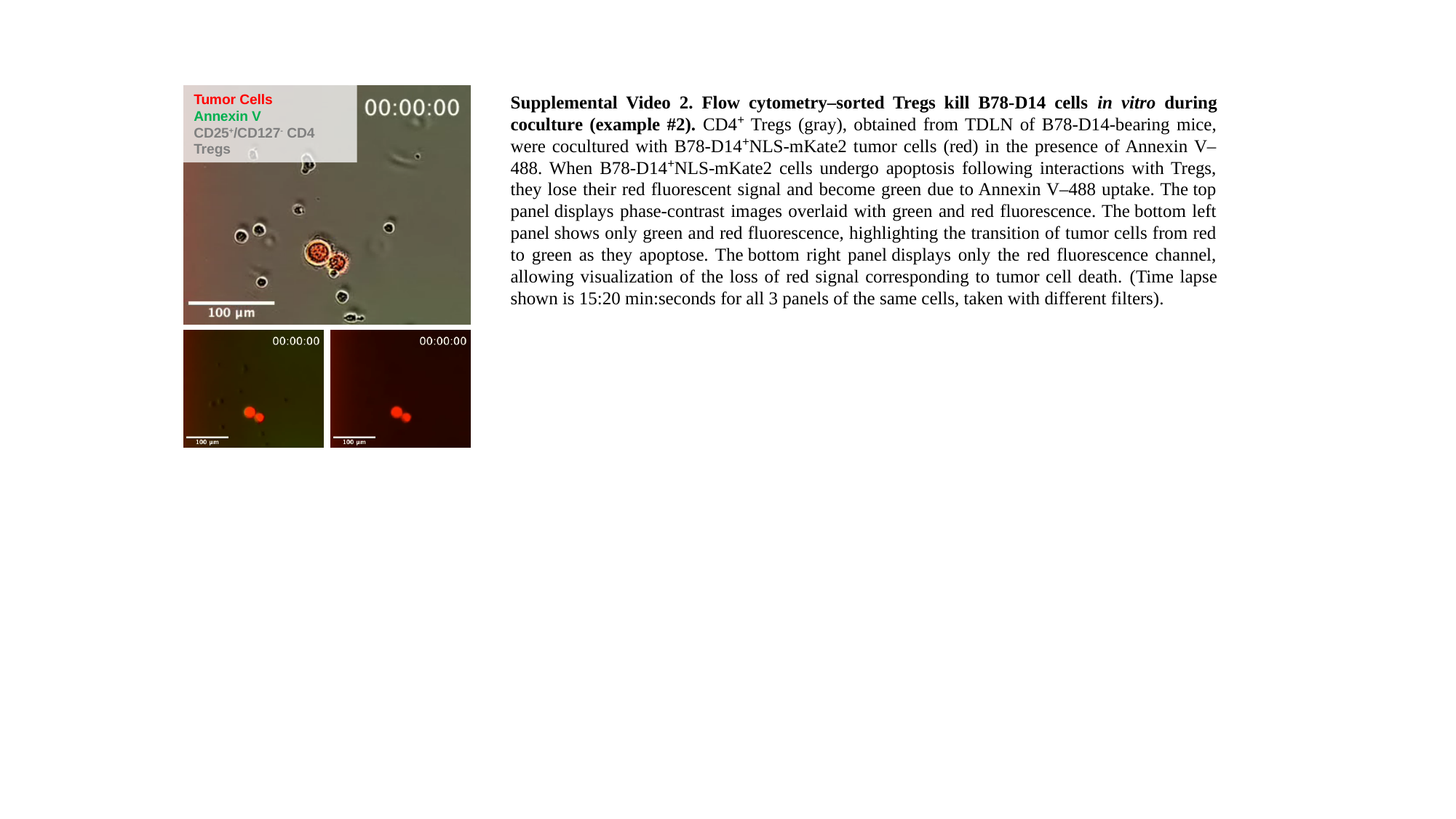

Tumor Cells
Annexin V
CD25^+^/CD127^-^ CD4 Tregs
Supplemental Video 2. Flow cytometry–sorted Tregs kill B78-D14 cells in vitro during coculture (example #2). CD4⁺ Tregs (gray), obtained from TDLN of B78-D14-bearing mice, were cocultured with B78-D14⁺NLS-mKate2 tumor cells (red) in the presence of Annexin V–488. When B78-D14⁺NLS-mKate2 cells undergo apoptosis following interactions with Tregs, they lose their red fluorescent signal and become green due to Annexin V–488 uptake. The top panel displays phase-contrast images overlaid with green and red fluorescence. The bottom left panel shows only green and red fluorescence, highlighting the transition of tumor cells from red to green as they apoptose. The bottom right panel displays only the red fluorescence channel, allowing visualization of the loss of red signal corresponding to tumor cell death. (Time lapse shown is 15:20 min:seconds for all 3 panels of the same cells, taken with different filters).
